# A Specialized Epithelial Cell Type Regulating Mucosal Immunity and Driving Human Crohn’s Disease

**DOI:** 10.1101/2023.09.30.560293

**Authors:** Jia Li, Alan J. Simmons, Sophie Chiron, Marisol A. Ramirez-Solano, Naila Tasneem, Harsimran Kaur, Yanwen Xu, Frank Revetta, Paige N. Vega, Shunxing Bao, Can Cui, Regina N. Tyree, Larry W. Raber, Anna N. Conner, Dawn B. Beaulieu, Robin L. Dalal, Sara N. Horst, Baldeep S. Pabla, Yuankai Huo, Bennett A. Landman, Joseph T. Roland, Elizabeth A. Scoville, David A. Schwartz, M. Kay Washington, Yu Shyr, Keith T. Wilson, Lori A. Coburn, Ken S. Lau, Qi Liu

## Abstract

Crohn’s disease (CD) is a complex chronic inflammatory disorder that may affect any part of gastrointestinal tract with extra-intestinal manifestations and associated immune dysregulation. To characterize heterogeneity in CD, we profiled single-cell transcriptomics of 170 samples from 65 CD patients and 18 non-inflammatory bowel disease (IBD) controls in both the terminal ileum (TI) and ascending colon (AC). Analysis of 202,359 cells identified a novel epithelial cell type in both TI and AC, featuring high expression of *<u>L</u>CN2*, *<u>N</u>OS2*, and *<u>D</u>UOX2*, and thus is named LND. LND cells, confirmed by high-resolution in-situ RNA imaging, were rarely found in non-IBD controls, but expanded significantly in active CD. Compared to other epithelial cells, genes defining LND cells were enriched in antimicrobial response and immunoregulation. Moreover, multiplexed protein imaging demonstrated that LND cell abundance was associated with immune infiltration. Cross-talk between LND and immune cells was explored by ligand-receptor interactions and further evidenced by their spatial colocalization. LND cells showed significant enrichment of expression specificity of IBD/CD susceptibility genes, revealing its role in immunopathogenesis of CD. Investigating lineage relationships of epithelial cells detected two LND cell subpopulations with different origins and developmental potential, early and late LND. The ratio of the late to early LND cells was related to anti-TNF response. These findings emphasize the pathogenic role of the specialized LND cell type in both Crohn’s ileitis and Crohn’s colitis and identify novel biomarkers associated with disease activity and treatment response.

## INTRODUCTION

Inflammatory Bowel Disease (IBD), comprising Crohn’s disease (CD) and ulcerative colitis (UC), is characterized by chronic, relapsing inflammation in the gastrointestinal tract (Kaser et al., 2010). CD can affect any portion of the gastrointestinal tract with inflammation that can span across all layers of the gut, while UC is localized to the colon and rectum and confined to the mucosa. IBD is believed to be driven from the complex interplay between environmental factors and genetic susceptibilities, resulting in dysregulated immune responses to environmental triggers and the breakdown of the epithelial barrier and intestinal homeostasis (Torres et al., 2017; Van Heel et al., 2001; Yang and Jostins-Dean, 2022). Genome-wide association studies have revealed more than 200 IBD-susceptiblility genes, which are involved in microbial sensing, antigen presentation, autophagy, T-cell signaling, and other immune-related pathways (de Lange et al., 2017; Jostins et al., 2012; Liu et al., 2015; Momozawa et al., 2018).

A wide range of cell types orchestrate intestinal host defense to environmental exposures. Characterizing cellular organizations and their rewiring in intestinal development and response to inflammation is of great importance to understanding IBD pathogenesis and to reveal novel potential treatment options. Recent studies utilized single cell and spatial omics profiling to provide an unbiased census of cell lineages and to characterize their functional states in healthy control and IBD samples (Boland et al., 2020; Bomidi et al., 2021; Burclaff et al., 2022; Elmentaite et al., 2021; Elmentaite et al., 2020; Fawkner-Corbett et al., 2021; Friedrich et al., 2021; Haber et al., 2017; Jaeger et al., 2021; Kanke et al., 2022; Li et al., 2021; Martin et al., 2019; Mitsialis et al., 2020; Moor et al., 2018; Parikh et al., 2019; Rosati et al., 2022; Smillie et al., 2019; Uniken Venema et al., 2019; Zhou et al., 2022). These studies successfully identified novel cells, dissected well-known cell types at high resolution, and revealed spatial, temporal, and functional heterogeneity of cellular compositions (Burclaff et al., 2022; Elmentaite et al., 2021; Fawkner-Corbett et al., 2021; Haber et al., 2017; Moor et al., 2018; Parikh et al., 2019). Comparing cellular differences between IBD and healthy controls, they found molecular and cellular alterations in disease, identified cellular modules associated with drug response, and built transcriptional links between the developing gut and childhood CD (Elmentaite et al., 2020; Friedrich et al., 2021; Martin et al., 2019; Smillie et al., 2019). While their main findings centered around immune cell signatures and immune-stromal interactions, few studies have shed light on epithelial cells regulating immune response and driving disease. Moreover, CD often involves the ileum where Paneth cells are abundant, while the colon contains no or few Paneth cells. Whether Crohn’s ileitis and Crohn’s colitis/UC act through a common mechanism remains largely unknown.

In this study, we combined bulk and single-cell RNA profiling, multiplexed imaging of proteins and RNAs, and spatial transcriptomics to study molecular and cellular remodeling and reorganization in active/inactive CD compared to non-IBD controls in both the terminal ileum (TI) and ascending colon (AC). We not only discovered rewiring of epithelial, stromal, and immune cells, but also identified a specific epithelial cell type in CD, which we named “LND cells” given their high expression of *<u>L</u>CN2*, *<u>N</u>OS2*, and *<u>D</u>UOX2*. LND cells are present in both the TI and AC of CD patients and expand with disease activity. LND cells specialize in regulating defense responses by recruiting and activating immune clells as signaling senders. Multiplexed imaging and spatial transcriptomics further demonstrated cross-talk between LND and immune cells. LND cells are a disease-critical cell type indicated by highly specific expression of IBD susceptibility genes. The presence of the LND cell type in both the TI and AC in CD patients suggests a common link to dysregulated host-environment interactions. A high resolution view of LND cells identified two subpopulations with different stem-potential and their ratio was associated with anti-TNF treatment response.

## RESULTS

### Cellular landscape of terminal ileum and ascending colon in non-IBD control and CD

We profiled 82 TI and 88 AC specimens from either endoscopic biopsies or surgical resection specimens across 83 unique individuals (65 CD patients and 18 non-IBD controls) using single-cell RNA-sequencing (scRNA-seq), representing one of the largest cohorts of CD patients profiled (Fig. 1A and Table S1). Non-IBD endoscopic specimens were collected from individuals presenting for colonoscopy for colorectal cancer screening or colon polyp surveillance without evidence of intestinal inflammation, while non-IBD surgical specimens were taken from normal adjacent tissue from patients undergoing surgical resection of endoscopically unresectable polyps in the cecum or ascending colon. Patient characteristics were as follows: ethnic background (CD: 77% white, 15% African American, 5% Asian, and 3% Hispanic; Control: 83% white, 11% African American, and 6% Hispanic), sex (CD: 63% female, 37% male; Control: 61% female, 39% male), and age (CD: 18-75; Control: 45-70) (Table S1). 6% of CD patients were treatment naïve, with the rest currently undergoing various treatments or previously treated for their CD symptoms (Table S1). Disease severity of each specimen was classified as active CD (31 mild, 9 moderate, and 11 severe) and inactive CD (58 normal and 17 quiescent) based on histopathologic analysis (Fig. 1B). The non-IBD specimens comprised 20 TI and 24 AC (Fig. 1B). In 77% of cases, matching TI and AC samples were collected from the same individual (Table S1).

**Figure 1.**
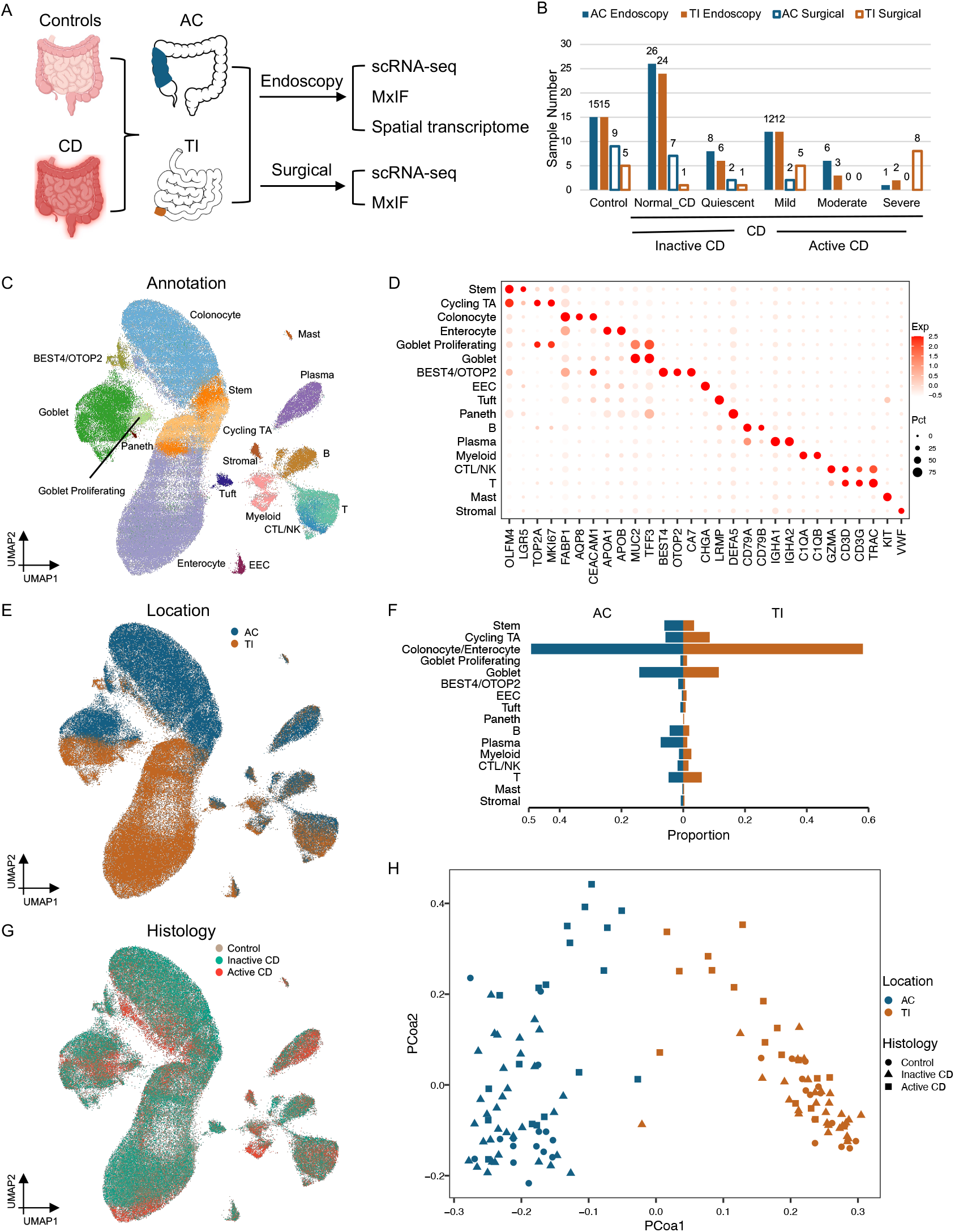
A) Schematic for processing endoscopic and surgical specimens from TI and AC for non-IBD controls, inactive and active CD patients. B) Summary of the number of samples in each group. C) UMAP of 155,093 cells from endoscopy samples colored by cell types. D) Dotplot showing markers for each cell type. E) UMAP of 155,093 cells colored by tissue origin, TI (brown) or AC (blue). F) Proportion of each cell type in TI (brown) and AC samples (blue). G) UMAP of 155,093 cells colored by disease status, control (tan), inactive (green) or active CD (red). H) MDS plot of cellular compositional differences across all endoscopic specimens.

After quality control, 155,093 cells from endoscopic specimens and 47,266 cells from surgical specimens were retained, and these datasets were analyzed separately (Table S2). Louvain clustering on single cells from endoscopic specimens revealed major cell types within the epithelial compartment (enterocytes/colonocytes, transit amplifying (TA) cells, stem cells, goblet cells, goblet proliferating cells, BEST4/OTOP2 cells, tuft cells, enteroendocrine cells (EEC), and Paneth cells - only in the TI), and the non-epithelial compartment (T cells, B cells, plasma cells, myeloid cells, cytotoxic T/natural killer (CTL/NK) cells, mast cells, and stromal cells) (Fig. 1C). Cell types were manually curated by known marker genes with TI enterocytes expressing *APOA1/APOB*, AC colonocytes expressing *AQP8*, TA cells expressing *MKI67*, stem cells expressing *LGR5*, goblet cells expressing *MUC2* and *TFF3*, goblet profilerating cells expressing both *MUC2* and *MKI67*, crypt absorptive cells expressing *BEST4* and *OTOP2*, Paneth cells expressing defensins such as *DEFA5* and *DEFA6*, T cells expressing *CD3D* and *CD3G*, B cells expressing *CD79A* and *CD79B*, plasma cells expressing *IGHA1* and *IGHA2*, myeloid cells expressing *C1QA* and *C1QB*, mast cells expressing *KIT*, CTL/NK cells expressing *GZMA*, and stromal cells expressing VWF (Fig. 1D). These results are consistent with cell types identified by previous scRNA-seq studies of the healthy human small intestine and colon (Burclaff et al., 2022; Fawkner-Corbett et al., 2021).

The largest transcriptomic difference was derived from absorptive cells from TI or AC (Fig. 1E). The most distinguishing marker genes for TI enterocytes were *APOA1* and *APOB*, which encode apolipoproteins and play a key role in intestinal lipid absorption. In contrast, AC colonocytes were marked by *AQP8*, a major transcellular water transporter (Fig. 1D). Differential transcriptional programs between TI enterocytes and AC colonocytes were identified (Fig. S1A), which reflected tissue-specific functions. Genes highly expressed in the TI were enriched in programs of protein/fat/vitamin absorption and processing, while genes highly expressed in the AC were enriched in nitrogen/sulfur metabolism (Fig. S1B), consistent with the nutrient digestion function of the small intestine and the interaction of the colon with microbial metabolites. Furthermore, signatures of region-specific development programs were also different between TI and AC. AC cells expressed hindgut-specific transcription factors such as *SATB2* and *CDX2* (Gu et al., 2022; Munera et al., 2017; Stringer et al., 2012), as well as the colon-specific chloride anion exchanger *SLC26A3*, also known as DRA (Down-regulated in adenoma)(Chatterjee et al., 2017) (Fig. S1A). In contrast, TI cells exhibit increased expression of machinery that organizes the brush border, including *VIL1* (Villin) and *CDHR2*, also known as protocadherin-24 (Crawley et al., 2014) (Fig. S1A).

Differences in goblet, EEC, and tuft cells between human TI and AC have not been fully investigated previously. We found that expression of region-specific genes identified in absorptive cells were also generally different in goblet, EEC, and tuft cells between TI and AC (Fig. S1C). Interestingly, there are region-specific genes that are also cell type-specific (Fig. S1C). For instance, *BEST2* was exclusively expressed in AC goblet cells, but not in TI goblet cells or other AC epithelial cell types (Figs. S1C and S1D) (Ito et al., 2013). *XBP1*, an ER stress response factor commonly expressed highly in secretory cells, was more highly expressed in TI goblet cells (Figs. S1C and S1D). SNPs in *XBP1* have been found to confer increased risk for developing IBD (Kaser et al., 2008), pointing towards an overload of protein production and ER stress that can potentially lead to secretory cell and barrier dysfunction, tipping the balance toward IBD. *RGS13* was exclusively expressed in TI tuft cells (Figs. S1C and S1D). *NTS* and *CLU* were exclusively expressed in TI EEC cells (Figs. S1C and S1D). EEC cells have been shown to act as facultative stem cells upon certain types of damage (Vega et al., 2019). Clusterin expression in TI EEC cells may implicate stem cell activation in the context of TI epithelial restitution as a response to damage, as clusterin was recently identified as a marker of revival stem cells (Ayyaz et al., 2019). Regarding immunological genes, AC goblet cells expressed more *IL1R2*, suggesting they were more sensitive to inflammatory stimuli (Fig. S1C). TI enterocytes expressed more *CCL25*, which is a chemokine ligand for CCR9 expressed on T cells responsible for inflammatory T cell recruitment (Wendt and Keshav, 2015) (Fig. S1A). While the TI and AC have distinct epithelial transcriptional programs, expression profiles of immune and stromal cells between the two sites largely overlapped, with similar cell types identified (Figs. 1C and 1E). In addition, the TI and AC exhibited similar composition of epithelial, stromal, and immune cells (Fig. 1F).

Transcriptomic differences were also observed as a function of disease activity (Fig. 1G). UMAP co-embedding revealed a shift of epithelial cell transcriptional state from non-IBD control, inactive, to active CD (Fig. 1G), indicating transcriptional changes driven by inflammation. Quantifying cellular compositional distances between individuals by scUniFrac (Liu et al., 2018) revealed that disease status is one of the main factors driving compositional shifts (Fig. 1H). Surprisingly, UMAP co-embedding revealed intermixing of cells from different individuals with the same disease status, suggesting only subtle if any patient-specific variability (Fig. 1G).

Analysis of 47,266 cells from surgical specimens generated similar results, with similar epithelial and non-epithelial cell types identified by marker genes and gene signatures (Figs. S2A, S2E, and S2G). The same differences between transcriptional programs of the TI and AC were observed in data generated from biopsies (Fig. 1E) and surgical resections (Fig. S2B). Furthermore, shifts in cell populations as a function of disease activity were also observed in the two specimen types (Fig. S2C and S2F). However, cell distributions between biopsies and surgical resections were different, with endoscopic biopsies dominated by epithelial cells and surgical specimens enriched for immune and stromal cells (Fig. S2D). This result reflects the superficial mucosal sampling of biopsies versus deeper submucosal sampling of surgical resections. For the rest of the analysis, we mainly focused on endoscopic specimens, which had a larger sample size with more evenly distributed disease status compared with surgical specimens.

### Distinct immune and stromal cellular organizations in active CD

Because CD is inherently characterized as an inflammatory disease, we set out to delineate changes within the immune and stromal compartments as a function of disease activity within our CD specimens. High resolution clustering revealed 12 populations of immune and stromal cells (Fig. 2A), with T cells expressing *CD3D* and *CD3G*, CTL/NK cells expressing *GZMB*, *KLRD1 (CD94),* and *NGK7*, B cells expressing *CD19* and *MS4A1 (CD20)*, and plasma cells expressing immunoglobins such as *IGHA1*, *IGHA2*, and *JCHAIN*. Proliferative T and B cells were also marked by a proliferative signature including *TOP2A* and *MKI67*. Within myeloid cells, mast cells were annotated by high expression of *KIT* and *FCER1A*, and neutrophils were marked by high expression of *S100A8*, *S100A9*, and *CXCR2* (Fig. 2B). We also recovered two types of macrophages. Resident macrophages expressed tissue residency markers *MRC1* and complement genes *C1QA* and *C1QB*, while recruited macrophages highly expressed inflammatory molecules, such as *NFKBIA*, *NFKBI*, *CXCL16*, and *CXCL9* (Fig. 2B). We identified fibroblasts expressing *PDGFRA* and ECM genes, such as *COL1A1*, *COL1A2*, and *COL6A1* and endothelial cells expressing *VWF* and *PECAM1* (Fig. 2B).

**Figure 2.**
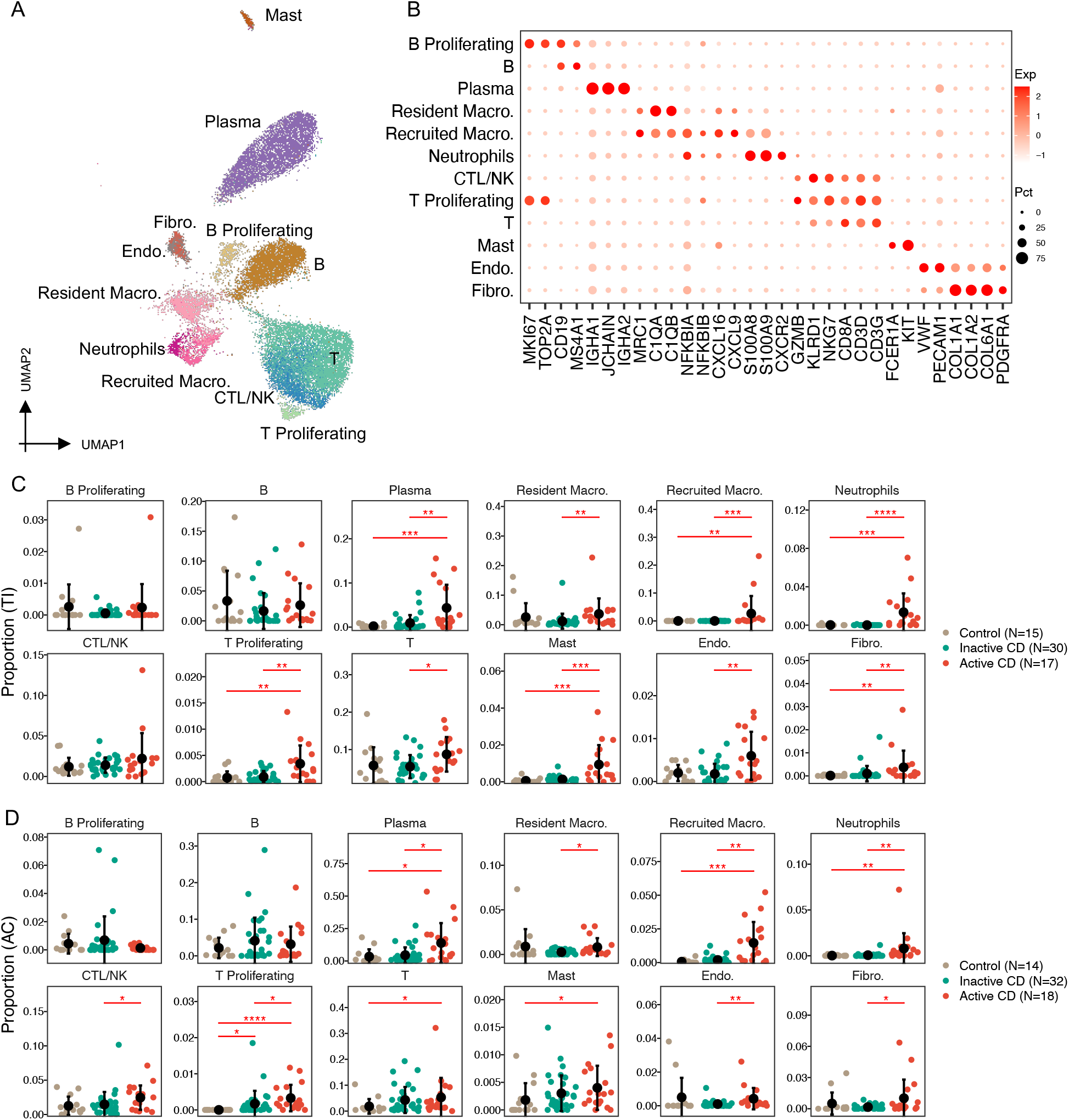
A) UMAP of immune and stromal cells, colored by cell type. B) Dotplot of markers in each cell type. C) Proportional changes of each immune and stromal cell type from non-IBD controls to inactive and active CD patients in TI. D) Proportional changes of each immune and stromal cell type from non-IBD controls to inactive and active CD patients in AC (* FDR<0.05, ** FDR<0.01, *** FDR<0.001, **** FDR<0.0001).

As expected, cellular composition within the immune and stromal compartments differed significantly between inactive and active CD (Table S3). Almost all immune and stromal cell numbers increased significantly in active compared to inactive CD (Figs. 2C and 2D), with the exception of B cells, proliferating B cells, and CTL/NK cells in the TI (Fig. 2C), and mast cells, B cells, proliferating B cells, and T cells in the AC (Fig. 2D). Among them, neutrophils and recruited macrophages showed the most significant elevation in active CD, in both TI and AC. The proportion of neutrophils increased from 0% to 1.3% (FDR=2e-6) in TI and from 0.06% to 0.7% in AC (FDR=0.002), which is not surprising as the presence of neutrophils is a hallmark of histologically active disease, while the proportion of recruited macrophages increased from 0% to 2.7% in TI (FDR=1e-4) and from 0.19% to 1.5% in AC (FDR=0.002)(Figs. 2C and 2D). Beyond cellular compositional changes, transcriptional upregulation of pro-inflammatory genes was observed in active CD compared to inactive CD within each cell type (Fig. S3A). Inflammatory signaling programs characteristic of the type 1 immune response appear to be upregulated in almost all immune cells; these include components of the JAK/STAT (*JAK3*, *STAT1*, *SOCS1*) pathway and a large variety of interferon regulatory factors (IRFs) and interferon stimulated genes (ISGs). Consequently, antigen-presentation genes, such as *CD74*, *B2M*, *HLA-A*, *HLA-B*, and others, were upregulated in both lymphoid and myeloid cell types (Fig. S3A). T cells were characterized by increases in both cytotoxic (*GZMB*, *NKG7*) and exhaustion phenotypes (*LAG3*, *TIGIT*) (Fig. S3A). Resident macrophages in active CD specifically increased expression of various cathepsins (such as *CTSB* and *CTSC*) that are important in microbial defense, as well as *CXCL9* that is important in further recruitment of CXCR3+ T cells (Figs. S3A).

Immune and stromal cell compositions were mostly unchanged between inactive CD and controls, with the exception of T proliferating cells in the AC, which was slightly increased in inactive CD (from 0% to 0.17%, FDR=0.02) (Fig. 2D). However, transcriptional upregulation of proinflammatory genes was still observed in a cell type-specific manner, such as overexpression of MHCII genes,such as *HLA-DRB5* and *HLA-DPB1* in B cells, and *HLA-DQB1* in both B cells andresident macrophages (Fig. S3B).

Analysis of surgical samples revealed similar cell types (Fig. S4A). Furthermore, we identified three subtypes of fibroblasts, *PDGFRA+* fibroblasts, *ABCA8+* fibroblasts, and *PDPN+* fibroblasts in both the TI and AC (Fig. S4A). The three subtypes had distinct expression profiles. *ABCA8+* fibroblasts had high expression of *ABCA8*, *CFD* and *SFRP2*, *PDGFRA+* fibroblasts had enriched expression of *PDGFRA*, *CXCL14*, and *ADGRL3*, and *PDPN+* fibroblasts were marked by high expression of *PDPN*, *MMP1*, and *SOD2* (Fig. S4B). The *PDGFRA+* fibroblasts in the surgical samples were similar to fibroblasts identified in TI and AC endoscopic biopsies (Fig. S4C). The *PDPN+* fibroblasts, also called activated or inflammatory fibroblasts, reside in the submucosa of the inflamed intestine outside of the lamina propria (Friedrich et al., 2021). They have been proposed as a central hub in IBD with an essential role in hematopoietic-stromal interactions (Friedrich et al., 2021; Martin et al., 2019; West et al., 2017). An increasing trend was observed for most stromal cell types in the TI (not implemented in the AC due to only two active CD surgical samples), especially *PDPN+* fibroblasts (Fig. S4D). Increases in recruited macrophages and neutrophils were observed in surgical specimens similar to the endoscopic biopsies (Fig. S4D). However, the increases were not statistically significant due to limited sample sizes in the TI surgical samples.

### A novel LND cell cluster specialized in regulating mucosal immunity

Deep annotation of epithelial cells revealed a continuum of cells consisting of stem, TA, early, intermediate, and mature enterocytes/colonocytes (Figs. 3A and 3C) (Burclaff et al., 2022; Fawkner-Corbett et al., 2021). Most interestingly, we identified a novel absorptive cell type that emerges and then expands during active inflammation in both the TI and AC. This subpopulation was marked by high expression of *LCN2*, *NOS2*, and *DUOX2*, therefore, we named it LND (Figs. 3B and 3D). LND cells were rare in non-IBD control tissues, increased marginally in inactive CD (0.3% to 1.3%, FDR=0.5 in TI; 0.06% to 0.6%, FDR=0.1 in AC), and expanded significantly in active CD (1.3% to 18.9%, FDR=1.7e-6 in TI; 0.6% to 17.8%, FDR=1e-5 in AC) (Figs. 3E and 3F; Table S3). The increase was observed both in the TI and AC, even though absorptive cells from both regions, as well as LND cells, have distinct transcriptomes. The proportion of LND cells was not associated with medication exposures (Fig. S5). LND cells increased at the expense of early, intermediate, and mature enterocytes/colonocytes, as well as BEST4/OTOP2 crypt-top cells as CD progresses from inactive to active disease (Figs. 3E and 3F; Table S3). The emergence and expansion of the LND cluster in actively inflamed tissues was also observed in surgical TI samples (Fig. S6).

**Figure 3.**
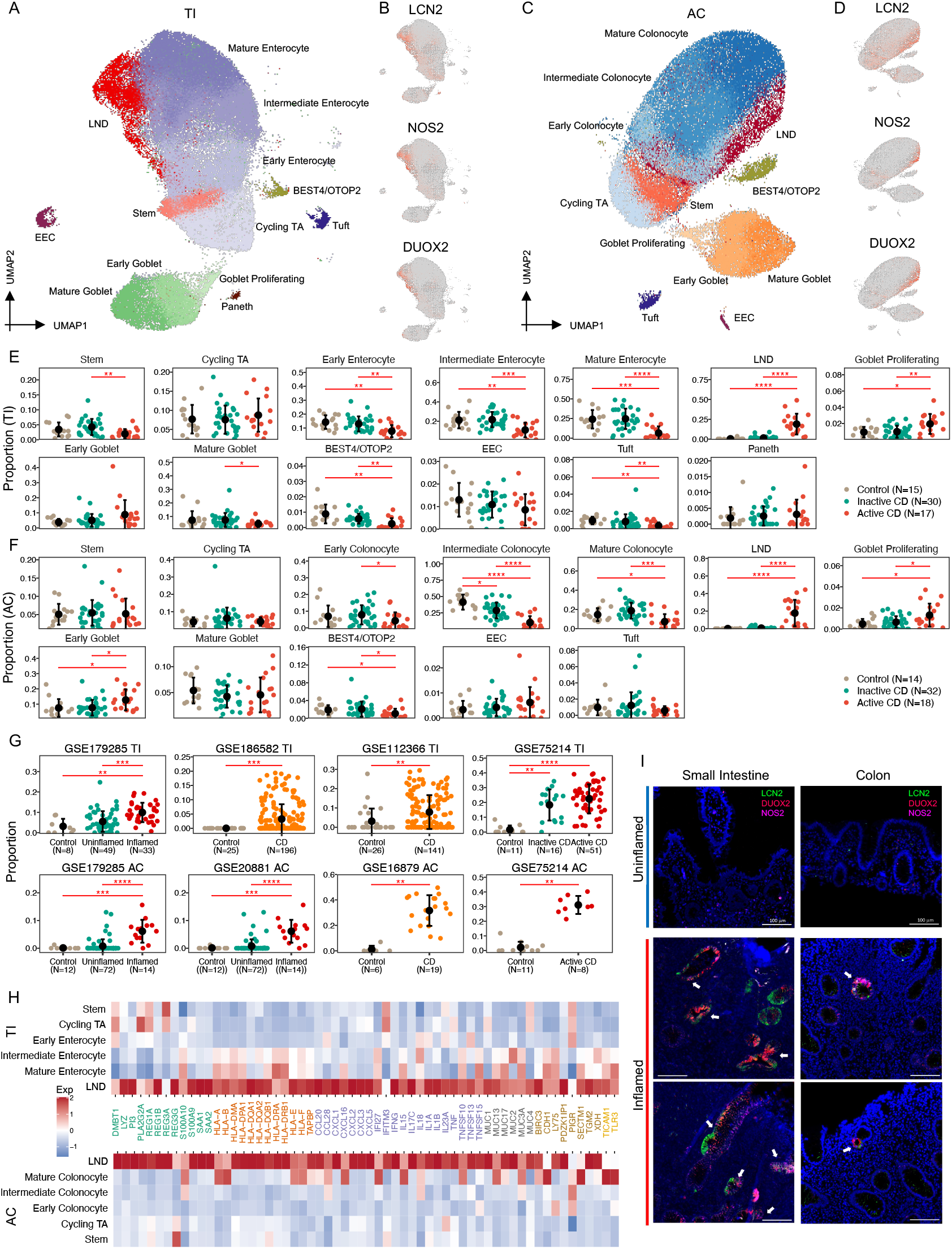
A) UMAP of 13 epithelial cell types in TI. B) UMAP labeled with expression of LCN2, NOS2, and DUOX2 in TI. C) UMAP of 12 epithelial cell types in AC. D) UMAP labeled with expression of LCN2, NOS2, and DUOX2 in AC. E) Proportional changes of each epithelial cell type from controls to inactive and active CD patients in TI. F) Proportional changes of each epithelial cell from controls to inactive and active CD patients in AC (* FDR<0.05, **FDR<0.01, ***FDR<0.001, **** FDR<0.0001). G) Proportional changes of LND cells with disease status in six independent cohorts (** FDR<0.01, ***FDR<0.001, **** FDR<0.0001). H) Heatmap of high expression of immune-related genes in the LND in both TI (top) and AC (bottom). I) HCR-FISH Co-staining of LCN2 (green), NOS2 (pink), and DUOX2 (red) on inflamed and uninflamed TI and AC tissues. The scale bar represents 100 µm.

We verified the emergence and expansion of LND cells in CD in six independent studies with publically available datasets and clinical information on disease activity. Consistently, LND cells were rare in healthy controls and emerged in uninflamed CD and expanded significantly in inflamed CD (Fig. 3G). Significant expansion of LND cells was observed from uninflamed to inflamed CD in both TI and AC in GSE179285 (FDR=1e-4 in TI and FDR=4e-9 in AC) (Fig. 3G). In GSE75214, LND cells also expanded from healthy controls to inactive CD in TI (FDR=4e-4) and to active CD in AC (FDR=2e-4) (Fig. 3G). A similar trend was detected in four other independent studies (GSE186582 and GSE12366 in TI, and GSE20881 and GSE66207 in AC) (Fig. 3G). The increase of LND cells at the expense of mature enterocytes/colonocytes was also observed in the six studies (Figs. S7 and S8).

Genes defining the LND cell cluster, *LCN2*, *NOS2*, and *DUOX2*, are all involved in host response to microbiota. LCN2, lipocalin 2, acts as an antimicrobial protein, which attenuates bacterial growth by binding and sequestering iron-scavenging siderophores (Goetz et al., 2002). LCN2 is a serum and fecal biomarker for intestinal inflammation (Stallhofer et al., 2015), and it has been reported to be increased in serum from CD patients (Scoville et al., 2019). NOS2, nitric oxide synthase 2, is an enzyme catalyzing the production of nitric oxide (NO), a broad-spectrum anti-bacterial agent (Muhl et al., 2011), and *NOS2* has been shown to be increased in colonic tissues from both CD and UC patients (Coburn et al., 2016). DUOX2, dual oxidase 2, produces hydrogen peroxide that supports mucosal oxidative antimicrobial defense. DUOX2 has been found to be upregulated in intestinal inflammation in a TLR-4-dependent manner (Burgueno et al., 2021) and is involved in NOD2-mediated antibacterial response (Lipinski et al., 2009). In addition to these marker genes, LND cells also express a high level of anti-microbial peptides (AMPs), including *DMBT1*, *REG1A*, *REG1B*, *REG3G*, *PI3*, *S100A9*, *LYZ*, *SAA1*, and *SAA2*, and upregulate transmembrane mucins (*MUC13*, *MUC17*, and *MUC3A*) that form the glycocalyx, which acts a physical barrier to luminal antigens (Fig. 3H). Downstream of immediate microbial defense, LND cells also overexpress genes that orchestrate immune responses. These include pattern recognition receptors *TLR3* and its interacting partner *TICAM1*, inflammatory signaling and immunity modulator *BIRC3*, antigen-presenting machinery (*HLA-B*, *HLA-A*, *HLA-DPA1*, *HLA-E*, *HLA-F*, *HLA-DQA1*, *HLA-DQB1*, *HLA-DQA2*, and *TAPBP*), and cytokines (*CCL20*, *CCL28*, *CXCL1*, *CXCL2*, *CXCL3*, *CXCL5*, *CXCL16*, *TNF*, *IFNG*, *IL13A*, and *IL17C*) (Fig. 3H). Functional analysis of genes upregulated only in LND cells compared to other epithelial cells found that they were enriched in antigen processing and presentation, Th17 cell differentiation, Th1 and Th2 cell differentiation, HIF-1 signaling pathway, and TNF signaling pathway (Fig. S9). These results suggest that the LND cell cluster that expands in active inflammation serves specialized functions of antimicrobial response and immunoregulation.

To confirm the presence and location of LND cells, we employed hybridization chain reaction-fluorescene in-situ hybridization (HCR-FISH) on inflamed and uninflamed TI and AC tissues to examine the co-occurrence of *LCN2*, *NOS2*, and *DUOX2* transcripts (Fig. 3I). In general, uninflamed tissues have stereotypical architectures with a well-delineated crypt/villus axis in the TI and regularly-spaced crypts in the AC. Inflammed tissues are generally characterized by villous and/or crypt loss and infilitration of immune cells, which distorts spatial architecture. *LCN2, NOS2*, and *DUOX2* expression was undetectable in uninflamed tissues (Fig. 3I). In both inflamed TI and AC, *LCN2, NOS2*, and *DUOX2* FISH staining co-localized in a subset of, but not all, epithelial cells, consistent with the scRNA-seq results (Fig. 3I). These results validate the presence of *LCN2*, *NOS2*, and *DUOX2* co-expressing cells and their induction in active CD.

### LND cells actively interact with immune cells

To identify the potential immunomodulatory function of the LND cells, we inferred cell-cell communications between LND and any other cell types using CellChat (Jin et al., 2021). We found that LND cells actively interact with immune cells as both signaling senders and receivers with similar patterns in the TI and AC (Figs. 4A and 4B). Compared with other epithelial cells (stem, cycling TA, enterocytes/colonocytes, tuft, BEST4/OTOP2, goblet, EEC), LND cells showed stronger cytokine-receptor interactions especially as signaling sources, suggesting a specialized role in regulating mucosal immunity (Fig. 4A and 4B). *CXCL2*, *CXCL3*, and *CXCL5,* which encode known neutrophil-attracting chemokines acting through CXCR1/CXCR2 receptors, were expressed significantly higher in LND cells (Figs. S10A and S10B). The LND-neutrophil interaction showed a similar interaction strength as the recruited macrophage-neutrophil interaction, implicating similar importance of this interaction in neutrophil recruitment and inflammatory response orchestration (TI: Fig. 4C; AC: Fig. S10D). LND was also the primary cell type for recruiting plasma cells through CCL28-CCR10 signaling (TI: Fig. 4C; AC: Fig. S10D), where *CCL28* was significantly expressed in LND cells and *CCR10* in plasma cells (Figs. S10A and S10B). The CCL28-CCR10 interaction is critical for regulating intestinal IgA production under homeostasis or infection (Hu et al., 2011). SAA1 is an acute phase protein in response to inflammation and tissue injury, which has strong chemotactic activity for neutrophils and macrophages mediated through FPR2 (Badolato et al., 1994; Dufton et al., 2010; Liang et al., 2000). *SAA1* was significantly upregulated in LND cells (Figs. S10A and S10B), leading to strong LND-neutrophil and LND-recruited macrophage interactions (TI in Fig. 4C; AC in Fig. S10D). We also detected LND-B/Plasma cells interactions through TNFSF13-TNFRSF13B (TI in Fig. 4C; AC in Fig. S10D), LND-T cells by NECTIN2-CD226, and LND-immune cells by CD55-ADERG5 (TI in Fig. S10C; AC in Fig. S10D). Furthermore, we identified interactions between LND and T cells via TNFSF15-TNFRSF25, LND-recruited macrophages via TNFSF10-TFNRSF10B and strong MHC-I signaling, but only in TI not in AC (Fig. S10C).

**Figure 4.**
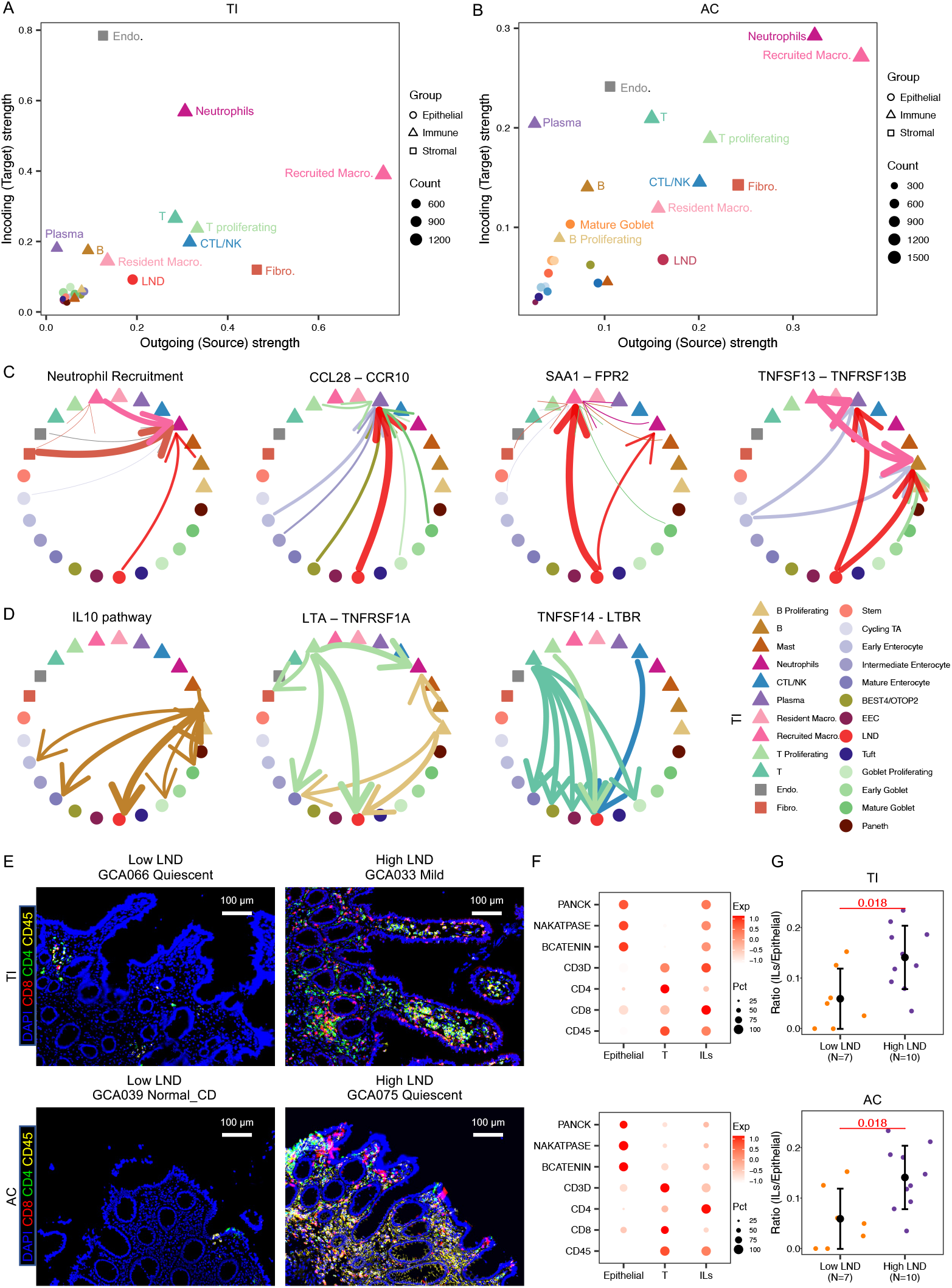
A) Scatterplot of incoming and outgoing interaction strength of each cell type in TI. B) Scatterplot of incoming and outgoing interaction strength of each cell type in AC. C) Circle plots show the intercellular communication for LND-recruiting neutrophils, LND-plasma interaction via CCL28-CCR10, LND recruiting macrophage and neutrophils through SAA1-FPR2, and LND-mast/plasma interaction by TNFSF13-TNFRSF13B. D) Circle plots show the intercellular signaling of LND cells targeted by a variety of immune cells including IL10, LTA-TNFRSF1A, and TNFSF14-LTBR. E) Multiplex images of CD8+, CD4+, and CD45+ cells in low and high LND in the TI (top) and AC (bottom). The scale bar represents 100 µm. F) DotPlot of marker genes in infiltrating lymocytes (ILs). G) The proportion differences of ILs between high and low LND patients.

LND cells not only actively participate in immune responses as signaling senders, but also respond to signals from immune cells as signaling receivers, although their activities as signal receivers were similar to other absorptive cells. A few examples of LND cells acting as signal receivers include B cell-LND by IL10 signaling and T/NK-LND by TNFSF14-LTBR in both TI and AC (TI in Fig. 4D; AC in Fig. S10D), and proliferative B/T cell-LND by LTA-TNFRSF1A only in TI (Fig. 4D).

Similar LND-immune interactions were observed for surgical TI samples (Fig. S11A). LND cells acted as both source and target for immune interactions. Consistent with the results found in endoscopic samples, LND cells were the primary cell type for recruiting plasma cells through CCL28-CCR10 signaling. Strong LND-neutrophil and LND-recruited macrophage interactions via SAA1-FPR2, LND-B/Plasma cells interactions through TNFSF13-TNFRSF13B, LND-T cells by NECTIN2-CD226, and LND-immune cells by CD55-ADERG5 were also detected (Fig. S11C). Furthermore, LND cells acted as signal receivers in the IL10 signaling and LTA-TNFRSF1A interaction (Fig. S11D). Interestingly, the PDPN+ fibroblast population serves as a strong signaling sender and is a main source for releasing CXCL2/CXCL3/CXCL5 in neutrophil recruitment (Figs. S11B and S11C).

We performed multiplexed protein imaging analysis on 55 tissues, of which 38 have single-cell RNAseq profiling (17 CD and 3 controls in TI, 15 CD and 3 controls in AC). We classified the multiplex imaging of CD patients into two categories based on the LND proportion reported in the single-cell RNAseq data, low and high LND. We found those with a high LND proportion had a significantly higher infiltration of lymphocytes into the epithelial submucosa in both the TI and AC compared to those with a low LND proportion (Figs. 4E and 4G). These infilitrating lymphocytes (ILs) were characterized by association of both epithelial (PANCK, NAKATPASE, and BCATENIN) and lymphocyte markers (CD3D, CD4, CD8, and CD45) (Fig. 4F). These results strengthen the association between CD activity, LND expansion, and immune cells infiltration. Since LND releases a variety of chemokines and cytokines (Fig. 3H) and actively interacts with immune cells (Figs 4A and 4B), they likely play a role in T cell recruitment and infiltration.

### LND is a CD/IBD-critical cell type

Genome-wide association studies (GWAS) on IBD have reported more than 200 genes involving 300 risk loci in multiple pathways (de Lange et al., 2017; Jostins et al., 2012; Liu et al., 2015; Momozawa et al., 2018). Previous studies applying these SNPs in a cell-type specific manner identified that these alterations in immune cells, especially T cells, are most strongly associated with IBD (Parikh et al., 2019). We combined GWAS-identified SNPs with single-cell RNA profiling to investigate the role of each cell type in CD. We utilized SNPsea (Slowikowski et al., 2014) to infer cell type-disease association by evaluating expression specificity of CD/IBD-associated risk genes in our scRNAseq data, with the assumption that risk genes specifically expressed in a cell type are likely driving disease by affecting a function unique to this cell type. Consistent with previous results (Parikh et al., 2019), we found T cells to be the most CD/IBD-associated cell type (FDR=0.001 in TI, FDR=9e-5 in AC), followed by recruited macrophages (FDR=0.001 in TI, FDR=0.03 in AC) and CTL/NK cells (FDR=0.02 in TI, FDR=0.03 in AC) in both TI and AC (Figs. 5A and 5B) (Table S4). T cell-disease association was driven by specific expression of *FYN*, *PTPRC*, *CD28*, *CD5*, *CD6*, *CARD11*, and other immune related genes (Fig. S12A). The macrophage-disease relationship was contributed to by specific expression of *LITAF*, *HCK*, *SLC11A1*, *MMP9*, *FCGR2A*, and *TNFAIP3*, and CTL/NK involvement was indicated by *KIF2DL4*, *IKZF3*, *TNFRSF18*, *CTSW*, and *PTPN22* (Fig. S12A).

**Figure 5.**
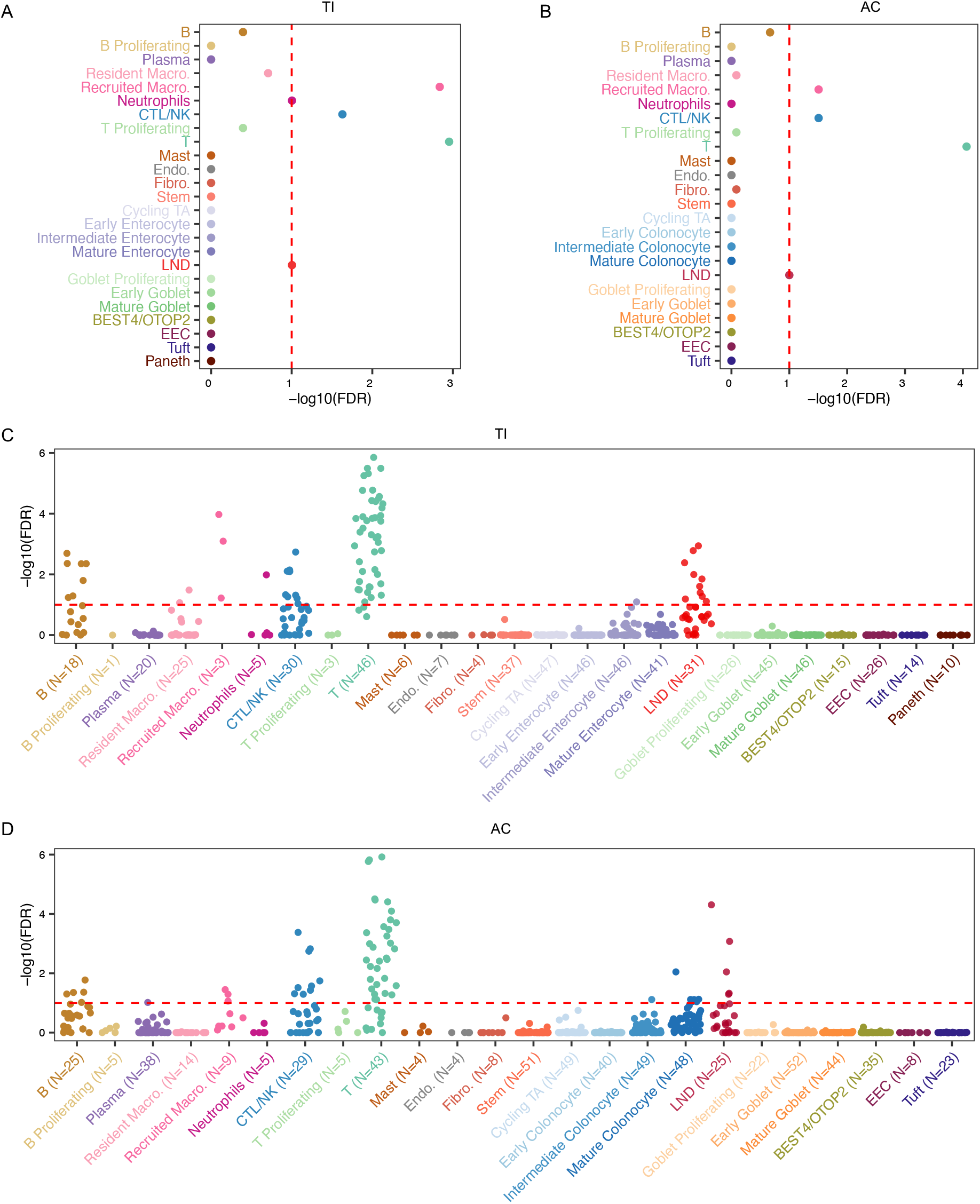
A) Significance of cell-type specific expression of IBD/CD-risk genes in TI. B) Significance of cell-type specific expression of IBD/CD-risk genes in AC. C) Significance of cell-type specific expression of IBD/CD-risk genes in each CD TI specimen. D) Significance of cell-type specific expression of IBD/CD-risk genes in each CD AC specimen.

Among epithelial and stromal cells, only LND cells were associated with CD/IBD, with marginal significance in both TI and AC (FDR=0.1) (Figs. 5A and 5B). *NOS2* was highly upregulated in LND compared to other cell types (Fig. S12A). A *NOS2* variant rs2297518 resulting in increased NO production has been associated with IBD (both CD and UC) (Dhillon et al., 2014). *CCL20* was also highly upregulated in LND cells (Fig. S12A), and one of its gene variants, rs111781203, has been reported to decrease the risk of IBD (Liu et al., 2015). Other CD/IBD-risk genes with high expression in LND cells were shown in Fig. S12A, such as *TNFRSF1A*, *STAT3*, *PLA2G2A*, *IRF1*, *TMBIM1*, and *PIGR*. Although genes highly expressed in LND cells, such as *DUOX2* and *LCN2*, have not been identified as CD/IBD-risk genes in large-scale GWAS studies, they have been reported to be associated with IBD risk or demonstrated to contribute to intestinal inflammation. Rare loss-of-function variants in *DUOX2* have been associated with increased plasma levels of IL-17C in patients, and *Duox2*-deficient mice confirmed increased IL-17C induction in the intestine (Grasberger et al., 2021). Biallelic mutations in *DUOX2* have been reported to be associated with very early-onset IBD (Kyodo et al., 2022). Depletion of LCN2 in mice leads to dysbiosis with increased intestinal inflammatory activity and an induction of Th17 cell differentiation (Kluber et al., 2021).

We observed extensive transcriptional heterogeneity of key genes in LND cells across CD patients. For example, *NOS2* was expressed highly in some patients, but its expression was low in others, although this gene was upregulated globally in the LND cluster (Fig. S12B). To address patient heterogeneity, we further evaluated disease association of each cell type on a per patient basis. We found T cells were significantly associated with CD in 43 out of 46 patients in TI, and 33 out of 43 patients in AC (FDR<=0.1). LND cells were significantly related to CD in 13 out of 31 patients in TI and 8 out of 25 patients in AC (FDR<=0.1) (Figs. 5C and 5D). In one CD patient (GCA062) with severe TI involvement, LND was significantly associated with CD, superceding the involvement of immune cells outside of recruited macrophages (FDR=0.001 for LND, FDR=0.001 for T cells, FDR=0.0008 for recruited macrophages). LND cells in this patient expressed high levels of *NOS2* and *CXCL5*, suggesting that this population is likely disease-critical (Fig. S12C). In contrast, no significant patient-specific association was observed for any other epithelial cell types, endothelial cells, or fibroblasts. These findings support the conclusion that LND cells might drive a significant portion of CD via dysregulated LND-immune cell communication.

### Developmental origins of LND cells

To infer the developmental origin of LND cells, we applied RNA velocity, an algorithm that predicts the future transcriptional states of each individual cell by the ratio of unspliced to spliced gene isoforms over the transcriptome (La Manno et al., 2018). As expected, we observed a cycling pattern for TA cells and a strong directional flow originating from stem cells, passing through early enterocyte, intermediate enterocyte, and ending in mature enterocytes in the TI (Fig. 6A). Interestingly, we found two origins that point toward LND cell development, one was from early enterocytes/stem cells, and the other was from mature/intermediate enterocytes, which paralleled the two subclusters of LND cells from high resolution clustering (Fig. 6B). The subcluster that originates from early enterocytes/stem cells was labeled “early LND”, while the other was labeled “late LND”. Partition-based graph abstraction (PAGA) analysis, which defines total connection strength between progenitor and differentiated cell populations (Wolf et al., 2019), also showed that early LND cells were associated strongly with early enterocytes, while late LND cells were linked to intermediate and mature enterocytes, as well as early LND cells (Fig. 6C). CytoTRACE analysis to infer the developmental potential of cell populations (Gulati et al., 2020) demonstrated that stem and TA cells had the highest inferred stemness score, followed by early enterocytes, early LND, intermediate enterocytes, late LND, and finally mature enterocytes (Fig. 6D). Our results indicate that LND cells can differentiate directly from stem/progenitor cells (early LND), or they may arise later (late LND) from intermediate/mature enterocyte or from early LND cells themselves.

**Figure 6.**
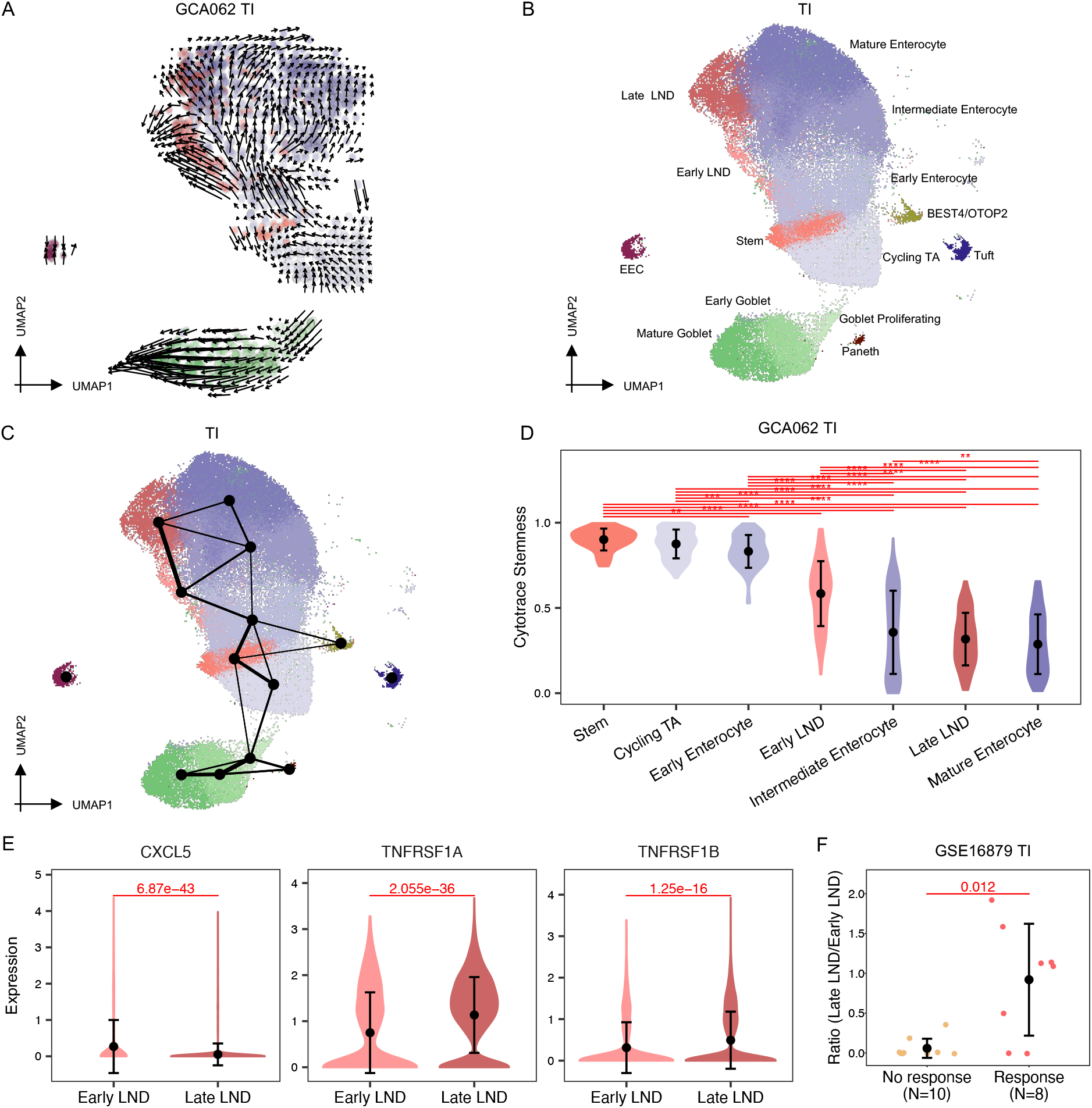
A) RNA velocity results mapped on the UMAP plot showing the predicted future transcriptional state of each cell. B) UMAP of early and late LND clusters in the TI. C) PAGA results mapped on the UMAP plot showing connectivity between cell types. D) Vilion plot comparing the developmental potential of each epithelial cell type predicted by CytoTRACE (** FDR<0.01, *** FDR<0.001, ****FDR<0.0001). E) Comparison of the expression of CXCL5, TNFRSF1A, and TNFRSF1B between early and late LND cells. F) Comparison of the ratio of late to early LND cells between anti-TNF responders and non-responders after the first dose of medication.

Differential expression analysis between early and late LND cells found that early LND cells are enriched in neutrophil chemoattractants (*CXCL3* and *CXCL5*), mucin (*MUC1* and *MUC4*), and anti-microbial genes (*DMBT1*, *PL2AG2A*, REG4, and *PIGR*), while late LND cells are enriched in lipid-metabolic genes (such as *APOC3*, *APOA4*, *APOB*, and *APOA1*), cytokines (*CCL20* and *CCL25*), *MUC3A, REG3G*, and TNF receptors (*TNFRSF1A*, *TNFRSF10B*, and *TNFRSF1B*) (Fig. 6E and Fig. S13A). They shared similar expression levels of *SAA1* and *CCL28*. Both early and late LND cells were increased as a function of disease activity, from normal non-IBD controls, inactive CD, to active CD (Fig. S13B). The ratio of early to late LND cells was also associated with disease activity (Fig. S13C), with early LND cells being enriched along the CD progression spectrum.

Since late LND cells expressed TNF receptors, we were curious whether the proportion of LND subclusters can predict anti-TNF response. We utilized the GSE16879 dataset (Arijs et al., 2009), which included 18 CD ileum patients assessed before and after their first anti-TNF treatment. In the Arijs et al. study, patients were classified as responders or non-responders based on endoscopic and histologic findings at 4-6 weeks after the initial treatment. We estimated the proportion of early and late LND cells by deconvolving bulk gene expression profiles through CIBERSORT (Newman et al., 2015). Although sample sizes were limited (n=10 responders and n=8 non-responders), we found that patients with higher proportions of late LND cells were more likely to respond to anti-TNF treatment (p=0.05). The proportion of early LND cells was not significantly associated with anti-TNF response. However, the ratio of late vs. early LND cells was significantly associated with anti-TNF response (p=0.012) (Fig. 6F).

### Colocalization between LND and immune cells

To investigate the organization and crosstalk between epithelial and immune cells, we used spatial transcriptomics to profile four CD samples selected for relatively high proportions of LND cells. The four samples consisted of two with active TI disease (GCA092 and GCA033) and two with active AC disease (GCA089 and GCA099) (Fig. 7A). As expected, LND marker genes, including *LCN2*, *NOS2*, *DUOX2*, and *CCL20/CCL28*, were coexpressed across spots in all four samples, indicating the existence of LND cells (Fig. S14). In contrast, expression of LND marker genes were not correlated with immune cell signatures, including *CD3D* for T cells, *CD8A* and *GZMB* for CD8+ T/NK, *MRC1* for resident macrophages, *NFKBIA* and *NFKBIB* for recruited macrophages, and *S100A8* for neutrophils (Fig. S14). Instead, high expression of LND marker genes in one spot was significantly correlated to high expression of immune cell signatures in its neighboring spots in all four samples using SpaGene (Liu et al., 2022) (Fig. 7B), suggesting heterotypic interaction between immune and epithelial cells (Fig. 7B). Specifically, *NOS2*, *LCN2*, and *DUOX2* all had a very significant colocalization with *GZMB*, *S100A8*, and *NFKBIA* in the GCA092_TI (FDR<3e-16). *DUOX2* colocalized with *CD8A* (FDR=2e-8) and *NOS2* with *NFKBIA* (FDR=2e-8), *CD8A* (FDR=2e-5), and *GZMB* (FDR=8e-5) in the GCA033_TI. *LCN2* colocalized with *NFKBIB* (FDR=3e-9) in the GCA089_AC. *NOS2* colocalized with *NFKBIA* (FDR=2e-9), and *LCN2* colocalized with *S100A8* (FDR=4e-8), *NFKBIA* (FDR=1e-5), and *NFKBIB* (FDR=2e-5) in the GCA099_AC. In comparsion, only marginally significant or insignificant colocalizations were found between the general epithelial genes (*KRT8* and *KRT18)* and immune markers in these samples (Fig. 7B).

**Figure 7.**
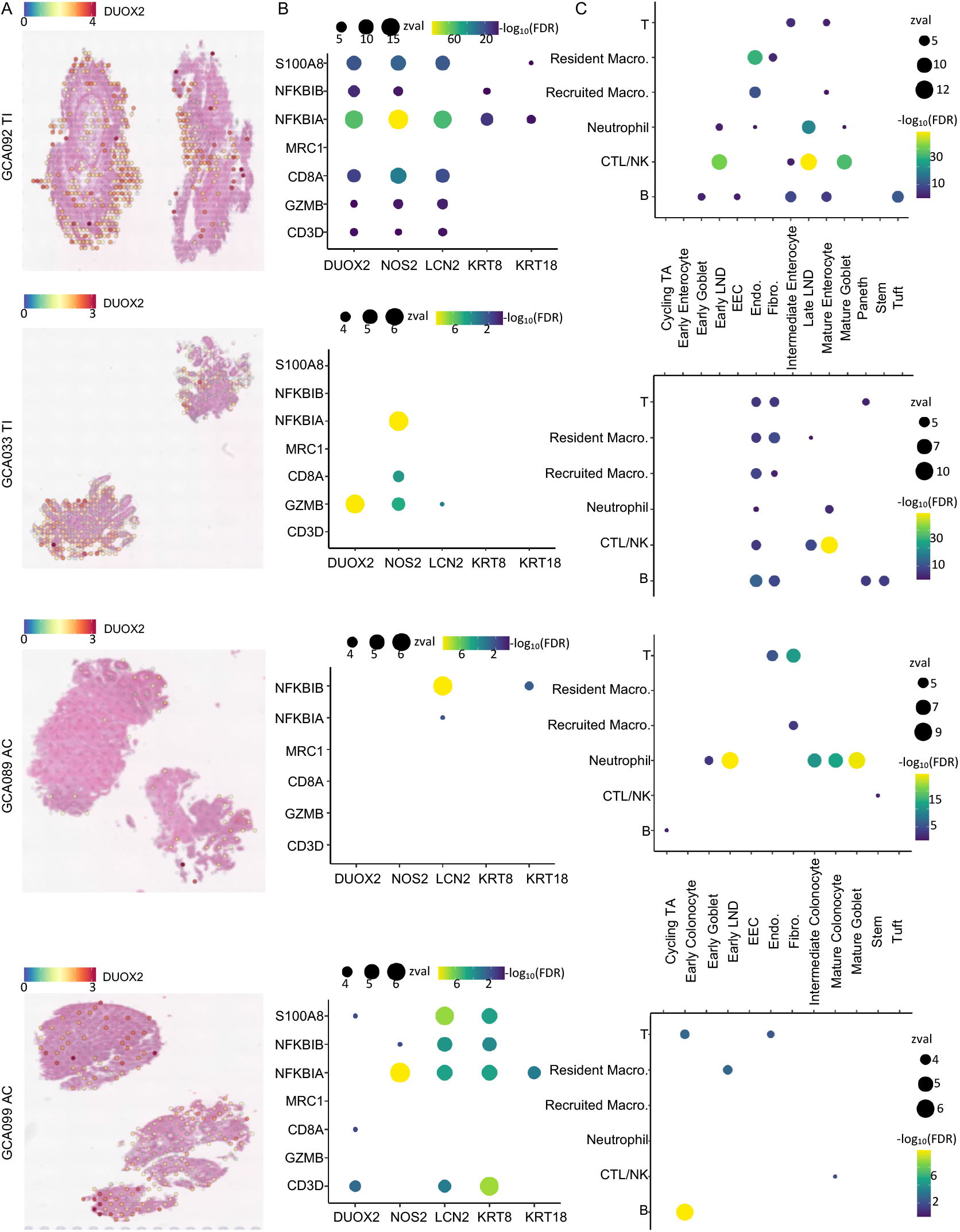
Spatial organization between LND and immune cells. A) H&E images for the four patient samples overlaid and colored by the expression of DUOX2. B) Dotplot of colocalization of LND markes (LCN2, NOS2, and DUOX2) and the general epithelial genes (KRT8 and KRT18) with immune signatures. Only significant colocalization (FDR<0.01) was included. Dot size denotes the z-value and color denotes the colocalization significance compared to random distribution. C) Dotplot of colocalization between epiethial and immune cells. Only significant colocalization (FDR<0.01) was included. Dot size denotes the z-value and color denotes the colocalization significance compared to random distribution.

To discover cell-type-specific spatial patterns and organization, we further deconvoluted spatial transcriptomics spots with cell-type profiles from scRNAseq using RCTD (Cable et al., 2022). We identified colocalization patterns between epithelial cells and immune cells based on decomposed cellular components by SpaGene (Liu et al., 2022) (Fig. 7C). In the GCA092_TI, the most signficiant association was found between late LND and CTL/NK (FDR=8e-35), followed by early LND-CTL/NK (FDR=2e-27) and late LND-Neutrophils (FDR=3e-16). In the GCA033_TI, significant association was observed between late LND-CTL/NK (FDR=3e-8). In the GCA089_AC, early LND and neutrophils were signficiantly colocalized (FDR=5e-24). In the GCA099_AC, LND and resident macrophages (FDR=1e-3) were significantly colocalized (Fig. 7C). In summary, LND cells were much more significantly associated with immune cells in all four inflamed specimens as compared to other epithelial cells, further demonstrating their specialized ability to interact with immune cells.

## DISCUSSION

In this study, we present a comprehensive single-cell atlas of 170 specimens from 83 individuals, consisting of 202,359 cells from the terminal ileum and ascending colon of human gut in non-IBD controls and inactive and active CD patients. We confirmed prior findings about region-specific transcriptomics to maintain physiologic function of the intestine and colon. Despite the distinct epithelial transcriptome between the TI and AC, we identified similar cellular rewiring in epithelial, immune, and stromal cell proportions with CD activity. For example, T cells, Mast, and recruited macrophages expand from inactive to active CD.

Most interestingly, we uncovered a new epithelial cell type, named LND, in both the TI and AC with high expression of *LCN2, NOS2,* and *DUOX2*. LND cells were rarely detected in non-IBD controls, but expanded significantly in active CD. Compared to other enterocytes/colonocytes, LND cells had high expression of anti-microbial proteins (such as *REG1A*, *REG1B*, *LYZ*, *PLA2G2A*, *SAA1*/*SAA2*), inflammatory cytokines (such as *CXCL2*, *CXCL3*, *CXCL5*, *CCL20*, *CCL28*), as well as antigen-presentation and processing genes, *STAT3* and *STAT1*, indicating a specialized immunoregulatory role. Cell-cell communication analysis supported that LND cells actively interact with a variety of immune cells as signaling senders. For example, LND cells release CXCL2, CXCL3 and CXCL5 to recruit neutrophils through CXCR2, and they can direct IgA+ plasma cell migration via CCL28-CCR10 interactions. Spatial transcriptomics further demonstrated the colocalization of LND cells and immune cells. The cross-talk between LND and immune cells highlights the role of LND in regulating mucosal immunity.

The intestinal epithelium is known to be the central coordinator of mucosal immunity, which requires a synergy of distinct epithelial cell types to promote homeostasis. These cell types carry out unique and specialized functions, including enterocytes/colonocytes for nutrient and water absorption, goblet cells for secreting mucins, Paneth cells for releasing antimicrobial peptides, and enteroendocrine cells for producing hormones. LND cells, in comparison, highly expressed some host defense-related genes which are cell-type specific in homeostatic conditions. For example, *REG1B, LYZ*, and *PLA2G2A*, which are antimicrobial peptides specifically released from Paneth cells, are highly expressed in LND cells. Consistently, previous studies found that expression of genes that are cell-type specific in homeostatic conditions was broadened across multiple cell types during infection (Haber et al., 2017). Therefore, LND cells are highly likely to be derived from enterocytes/colonocytes under chronic inflammatory stress, leading to specialized functions in immunoregulation. Studying the developmental origins of LND cells also supports that LND cells originate from early enterocytes or intermediate/mature enterocytes.

LND cells not only had high expression of cell-type specific genes as mentioned above, but also showed high expression of IBD/CD GWAS-risk genes, such as *NOS2*, *CCL20*, *TNFRSF1A,* and *STAT3*. The specific expression of IBD/CD-risk genes suggest LND cells are a critical disease cell type. The disease-association of LND cells was quite heterogenous across patients. In TI, LND cells in ∼ 30% of CD patients showed significant disease association and were ranked the second most important cell type. The heterogeneity of LND cells also reflects the complex and multifactorial pathogenesis of CD. In addition to IBD/CD-risk genes, LND cells were marked by high expression of additional genes previously demonstrated to modulate colitis, indicating their potential pathogenic role. Our studies identified that hematopoietic-LND cell interactions play an important role in regulating host response and driving CD, which extends previous findings emphasizing hematopoietic-stromal interactions as a central hub in IBD pathology (Friedrich et al., 2021; Martin et al., 2019; West et al., 2017).

Taken together, our study identified a novel LND cell populations with unique molecular features enriched in immunoregulation, providing a better understanding of the mechanisms sustaining the pathogenic process in Crohn’s Disesase. Our results indicate that LND marker genes and their cellular proportion could have clinical significance as markers of disease activity, risk for disease progression, or likelihood of anti-TNF response. Our findings establish the possibility of meet evolving clinical needs with characterization and personalized treatment of CD at the molecular level, which would greatly benefit future clinical studies.

## Materials and Methods

### Human specimen collection and processing

The study protocol was approved by the Institutional Review Board at Vanderbilt University Medical Center. Written informed consent was obtained from non-IBD control and CD subjects to obtain terminal ileum (TI) and ascending colon (AC) tissue at the time of scheduled endoscopic procedures. TI and AC tissues from non-IBD control and CD subjects undergoing surgical resection were also obtained from under a separate IRB protocol in coordination with the Comparative Human Tissue Network (CHTN). All samples were obtained as a part of the clinical trial “Combinatorial Single Cell Strategies for a Crohn’s Disease Gut Cell Atlas”, identifier NCT04113733 (clinicaltrials.gov).

Between December 2019 and January 2022, endoscopy subjects were prospectively recruited in the IBD clinic or GI endoscopy unit at Vanderbilt University Medical Center prior to colonoscopy for CD disease activity assessment or non-IBD indications including colorectal cancer screening or polyp surveillance. Surgical resection subjects were those undergoing resection for CD-related complications or other non-inflammatory indications, including endoscopically unresectable polyps. Patient participation in the current study ended after tissue samples were obtained. Exclusion criteria for the study were: pregnancy, known coagulopathy or bleeding disorders, known renal or hepatic impairment, history of organ transplantation, or inability to give informed consent. After appropriate exclusions, there were 65 CD subjects with varying disease activity and 18 non-IBD controls.

For all participants, demographics including age, gender, medical history, and medication use were determined from participant reporting and review of the electronic medical record. Tissue biopsies for research purposes in the TI and AC were obtained as follows: fresh tissue biopsies were placed in chelation buffer (4mM EDTA, 0.5 mM DTT in DPBS) for further processing and scRNAseq analysis, and an adjacent set of tissue biopsies were formalin-fixed and paraffin-embedded (FFPE) for research blocks. 5 µm sections were used from each FFPE block and stained with hematoxylin and eosin (H&E) and examined in a blinded manner by a gastrointestinal pathologist (MKW) and graded accordingly as: inactive (normal, quiescent) or active (mild, moderate, or severe activity). All associated study data were collected and managed using Research Electronic Data Capture (REDCap) electronic data capture tools hosted at Vanderbilt (Harris et al., 2019; Harris et al., 2009), including Clinical Data Interoperability Services, such as Clinical Data Pull (Cheng et al., 2021) and e-consent (Lawrence et al., 2020).

### Single-cell encapsulation and library generation

Single-cell RNA-seq data were generated from human biopsy and surgical specimen similarly to (Chen et al., 2021; Simmons and Lau, 2022). For surgical specimens that are considerably larger, a representative portion (∼2mm^2^) of the tissue was used. Briefly, tissues were incubated in a chelating buffer (0.5M EDTA, 0.1M DTT in DPBS) for 1.25hrs, and then transferred to cold active protease (5 mg/ml Protease from Bacillus licheniformis, 2.5 mg/mL DNase in PBS) for 25 minutes at 4°c. Tissues were then pipetted 10-20 times to yield single cells. Cell suspensions were filtered, washed, and inspected for count and quality before loading onto inDrops for microfluidic capture. inDrops scRNA-seq was performed as described (Banerjee et al., 2020; Klein et al., 2015). Single-cell libraries were prepared for sequencing as documented (Southard-Smith et al., 2020). Libraries (consisting of an estimated 2000-3000 cell transcriptomes) were sequenced at ∼125 million reads each on the Novaseq6000.

### HCR-FISH

HCR FISH was performed for three targets mRNAs using three DNA probe sets DUOX2, LCN2, and NOS2, using the HCR™ RNA-FISH Protocol for FFPE tissue sections (Choi et al., 2018). Tissue slides were baked at 60°C for 1 hour, followed by tissue deparaffinization by immersing slides in Xylenes, 3X for 5 minutes. After deparaffinization, slides were incubated in 100% Ethanol, 2X for 3 minutes. Rehydration of tissue slides was done by series of graded ETOH washes at 95%, 70% and 50% concentrations followed by nanopure water wash. After the rehydration steps, slides were immersed for 15 min in Tris-EDTA buffer (PH 9.0) at 95°C. Tris-EDTA buffer temperature was slowly cooled down to 45°C in 20 minutes, by adding nanopure water every 5 min. Slides were kept in nanopure water for 10 min at room temperature, followed by PBS1X wash. Proteinase K was introduced at 0.5 µL of 20 mg/1mL PBS1X concentration, for 10 min at 37°C, followed by PBS1X washes. 200 µL of Probe Hybridization Buffer was added on top of each tissue sample for pre-hybridization and slides were kept in humidified chamber, at 37°C, for 10 min. Probe solution was prepared by adding 0.4 µL of 1µM Stock/100 µL of probe hybridization buffer at 37°C for DUOX2 and LCN2 probe sets and 0.8 µL of 1µL of 1 µM stock/100 µL of probe hybridization buffer at 37°C for NOS2 probe set. Pre-hybridization solution was removed from tissue slides and 100 µL of the Probe solution was added on top of each tissue sample. Sample slides were covered with parafilm and incubated overnight at 37°C in the humidified chamber. Excess probes were washed by incubating slides at 37°C in: a) 75% of probe wash buffer/25% 5X SSCT for 15 min, b) 50% of probe wash buffer/50% 5X SSCT for 15 min, c) 25% of probe wash buffer/75% 5X SSCT for 15 min d)100% 5X SSCT for 15 min. Slides were immersed in 5X SSCT for 5 min at room temperature. For pre-amplification, 200 µL of amplification buffer was added on top of each tissue sample for 30 min at room temperature. 2 µL of 3 µM stock hairpins h1 and h2 (per slide), for each probe set, were separately heated at 95°C for 90 seconds and cooled to room temperature in the dark for 30 min. Hairpin solution was prepared by adding snap-cooled hairpins h1 and snap-cooled hairpins h2 to 100 µL of amplification buffer at room temperature. Pre-amplification buffer was removed and 100 µL of the hairpin solution was added on top of each tissue sample. Slides were incubated overnight ≥12h at Room temperature. Excess hairpins were removed by incubating slides in 5X SSCT at room temperature for 1 x 5 min, 2 × 15 min and lastly 1 X for 5 min. Slides were dried by blotting edges on a kimwipe. 100 µL of Hoechst stain (1:100 dilution) was added on top of each tissue slide and slides were incubated at room temp for 5 min. Cover slipping was done by using Invitrogen Prolong™ Gold antifade reagent. Slides were imaged using the Aperio Versa slide scanner (Leica). Probe sets were designed by Molecular Instruments: LCN2 Probe set: probe set size 13 targeting NM_005564.5, DUOX2 Probe set: probe set size 20 targeting NM_014080.5, NOS2 Probe set: probe set size 20 targeting NM_000625.4.

### Multiplex Immunofluorescence and Image Analysis

Multiplex immunofluorescence (MxIF) imaging was performed on FFPE sections at 4 µm after standard histological processing and antigen retrieval. Slides were iteratively stained using a fluorescence-inactivation protocol, as performed previously (Herring et al., 2018; Vega et al., 2022), using directly labeled antibodies incubated overnight at 4°C. Slides were scanned using the Aperio Versa (Leica) at 20X magnification, and then were photo-inactivated with an alkaline peroxide solution for repeated staining and imaging cycles until images for all analytes were acquired. A validated antibody panel was used, including DAPI, NAKATPASE, PANCK, CD8, CD4, CD45, etc.(Chen et al., 2021). Images were computationally registered and corrected for illumination and autofluorescence against periodic blank imaging rounds without antibody staining. Cells were segmented with an algorithm modified from one published (McKinley et al., 2022), using a combination of Ilastik machine learning, watershed using multiple membrane markers, and membrane completion. Cells meeting a certain quality thresholds of size were kept. The mean, standard deviation, median, and maximum staining intensity for each protein was quantified with respect to the whole cell, cell membrane, cytoplasm, and nucleus. Location, area, and shape metrics were obtained. Cells were clustered based on the similarity of protein intensity profiles and each cluster was annotated by positive expression of known marker genes.

### Spatial Transcriptomics

Spatial transcriptomics was performed using the Human FFPE Visium platform, as described previously (Heiser et al., 2023). FFPE sections (5 µm) of biospies were cut directly into 6.5mm × 6.5mm capture areas of Visium FFPE spatial gene expression slides (10X Genomics). Visium slides were temporarily coverslipped, stained with hematoxylin and eosin, and imaged in brightfield at 20X magnification using Y (Leica) prior to tissue permeabilization, probing, and library prep according to the Human Visium FFPE protocol (10X Genomics). Sample libraries were sequenced on the NovaSeq6000 sequencer (Illumina). Resulting sequencing data were aligned using 10X Genomics Space Ranger version 1.3.0 (10X Genomics).

### Single-cell RNAseq alignment and quality control

Single-cell RNAseq reads were filtered, demultiplexed, and quantified by dropEst (Petukhov et al., 2018) to generate cell-by-gene count matrices. Specifically, reads with expected structure were kept, and cell barcodes and UMI were extracted by dropTag. Demultiplexed reads were aligned to the human reference transcriptome GRCh38 using STAR (Dobin et al., 2013). Uniquely mapped reads were quantified into UMI-filtered counts by dropEst. Cells with >40% mitochondria reads, or <500 UMI counts, or <200 or >6,000 genes expressed were considered as low quality and excluded. After this rough quality control, each sample was manually checked to remove those clusters of empty droplets (low number of UMI and genes, and no distinct markers) and clusters of doublets (high number of UMI and genes, and markers from two different cell types). Samples with cells less than 100 were excluded from the downstream analysis. Outliers and batch effects were detected using scRNABatchQC (Liu et al., 2019).

### Single-cell RNAseq data analysis

Single-cell RNAseq count matrices were normalized to 10,000 and the top 2,000 highly variable genes were selected by fitting the variance-mean relationship in the Seurat package (Butler et al., 2018; Stuart et al., 2019). The normalized data were scaled to z-scores and principal component analysis was performed to reduce dimension. The top 10 principle compoents were used to generate the UMAP embedding for visualization and to to build the *k*-nearest neighbor graph (k=20). Louvain clustering at a resolution of 0.8 was applied on the graph to partition cells into non-overlapping groups by the Seurat. Cell clusters were automatically annotated by a marker-based approach scMRMA (Li et al., 2022) and were further manually curated using cluster-specific genes from the differential expression analysis. A divide-and-conquer strategy was adopted to provide a more precise clustering for those minor cell populations, such as immune cells and fibroblasts. Those minor cell populations identified from the initial clustering were extracted, reclustered, and reannotated. The clustering and annotation results served as the input to scUnifrac (Liu et al., 2018) to quantify cell compositional distances across samples, which considered both cellular compositions and similarities. Multidimensional scaling was used to map each sample into a space based on pairwise distances from scUniFrac.

### Cell type deconvolution for bulk transcriptomics data

CIBERSORT (Newman et al., 2015) was applied to characterize the cell composition of bulk RNAseq data using single-cell transcriptional profiles of each cell type from TI and AC as the reference. The signature matrix was created from average expression of the top 100 marker genes in each cell type. Default parameters were used to implement CIBERSORT, except that the parameter of quantile normalization of bulk mixture was set to False.

### Cell-cell interaction analysis

CellChat (Jin et al., 2021) was used to infer communications between cell types through ligand-receptor interaction analysis from single-cell RNAseq data of TI and AC separately. The standard workflow was followed with the normalized data and the annotated cell types as inputs. The built-in database CellChatDB.human involving 1,939 interactions was used as a reference to screen potential ligand-receptor interactions. The communication probablility was quantified between cell types having at least 10 cells. The average gene expression per cell type was caculated without trimming.

### Development trajectory analysis

RNA velocity (La Manno et al., 2018) was applied to infer lineage relationships between epithelial cell populations and predict future transcriptional state of a single cell. First, the loom file including spliced/unspliced matrices was generated from the bam file using Velocyto. Then velocity was calculated by the function RunVelocity in the SeuratWrapper package with default parameters. The velocity was plotted on the pre-computed UMAP embedding and colored by the annotated cell types.

CytoTRACE (Gulati et al., 2020) was performed to predict stemness status from single cell RNAseq data based on the assumption that the number of genes expressed in a cell decreases during differentiation. CytoTRACE was implemented with default parameters and the raw count matrix of each sample as the input. A CytoTRACE score was assigned to each cell based on its differentiation potential, with higher score indicating higher stemness. CytoTRACE scores from different samples were grouped by cell types and score differences between two cell types were compared by Wilcoxon rank-sum test.

Partition-based graph abstraction (PAGA) (Wolf et al., 2019) was used to reconstruct lineage relationships of epithelial cell populations. First, the Seurat object was converted to h5ad file for the PAGA input. Then, a neighborhood graph was computed based on the size of local neighborhood of 50 and the number of PCs of 30 using scanpy. Finally the connections between cell types were quantified. The connections of weight less than 0.2 were removed.

### Cell types associated with CD/IBD-risk Loci

SNPsea algorithm (Slowikowski et al., 2014) was used to identify cell types associated with CD/IBD-risk SNPs based on the assumption that genes specificity to a cell type is an indicator of its importance to the cell type function. Thus if one cell type have significant enrichment of specific genes associated with GWAS risk loci, this cell type is highly likely to be pathogenic and critical to the disease. The CD/IBD-risk SNPs were compiled from two GWAS studies (Jostins et al., 2012; Liu et al., 2015), which reported 344 loci in total. A pseudobulk dataset for each cell type in CD was generated by summing all UMI counts for each gene in each cell type and adding pseudocount of 1. The data were then normalized by DESeq2 to remove the effects introduced by cell cluster-sizes. SNPsea was run with defult parameters and all genes in a SNP’s linkage interval are accounted when calculating scores. The p-values were further adjusted by the Benjamini-Hochberg multiple testing procedure.

### Cell type deconvolution and spatial colocalization

Robust Cell-Type Decomposition (RCTD) (Cable et al., 2022) in the spacexr package was applied to deconvolve cell type compositions of each spot. Single-cell RNAseq data and cell types annotations from all TI and AC samples were used as the reference to decompose spatial TI and AC samples, respectively. The anchor-based integration workflow in the Seurat was also used to predict the underlying composition of cell types in each spot and similar results were obtained.

SpaGene (Liu et al., 2022) was used to quantify colocalization of markers of epithelial genes (*KRT8* and *KRT18*) and LND (*LCN2*, *NOS2*, and *DUOX2*) with immune cell signatures (*CD3D*, *CD8A*, *GZMB*, *MRC1*, *S100A8*, *NFKB1A*, and *NFKB1B*). Z-scores and FDR values were generated to estimate the significance of spatial connections of two genes (such as *NOS2* and *CD8A*) compared to random distributions. SpaGene was also performed to quantify colocalization between epithelial and immune cells based on the inferred composition of each cell type from RCTD (Cable et al., 2022). Z-scores and FDR values were produced to estimate the significance of spatial colocalizations of two cell types (such as LND and Neutrophils) compared to random connections.

### Acknowledgements

We thank Dr. Nicholas Zachos and I-Ling Chiang for their helpful contributions. This work is part of the Gut Cell Atlas Crohn’s Disease Consortium funded by The Leona M. and Harry B. Helmsley Charitable Trust and is supported by a grant from Helmsley to Vanderbilt University Medical Center (G-1903-03793) (http://www.helmsleytrust.org/gut-cell-atlas/). This work was also funded National Institutes of Health (NIH) grants (P01AI139449, R01DK103831, R01 DK128200), National Cancer Institute grants (NCI) (P50CA236733, P01CA229123, and U54 CA274367), Veterans Affairs Merit Review grants I01BX004366 (LAC), I01CX002171 (KTW), and I01CX002473 (KTW), Department of Defense PRCRP Impact Award W81XWH-21-1-0617 (KTW), Crohn’s & Colitis Foundation Senior Research Award 703003 (KTW), and NCI/NIH Cancer Center Support Grant P30CA068485. Additional support was provided by NIH grant P30DK058404 (Vanderbilt Digestive Disease Research Center) and NCATS/NIH grant UL1TR000445 (Vanderbilt Institute for Clinical and Translational Research). Whole slide imaging and quantification were performed in the Digital Histology Shared Resource at Vanderbilt University Medical Center. Surgical resection specimens were provided by the Cooperative Human Tissue Network (CHTN), which is funded by National Cancer Institute grant UM1CA183727.

## Supplementary Tables

Table S1. Metadata per sample, including sample ID, patient ID, endoscopy/surgical, specimen location, and disease status.

Table S2. Technical statistics per sample, including the number of cells and genes, and the total UMI.

Table S3. Cellular compositions in each sample.

Table S4. Cell type-disease association in each sample.

