## Supplemental figures for "A Specialized Epithelial Cell Type Regulating Mucosal Immunity and Driving Human Crohn’s Disease"

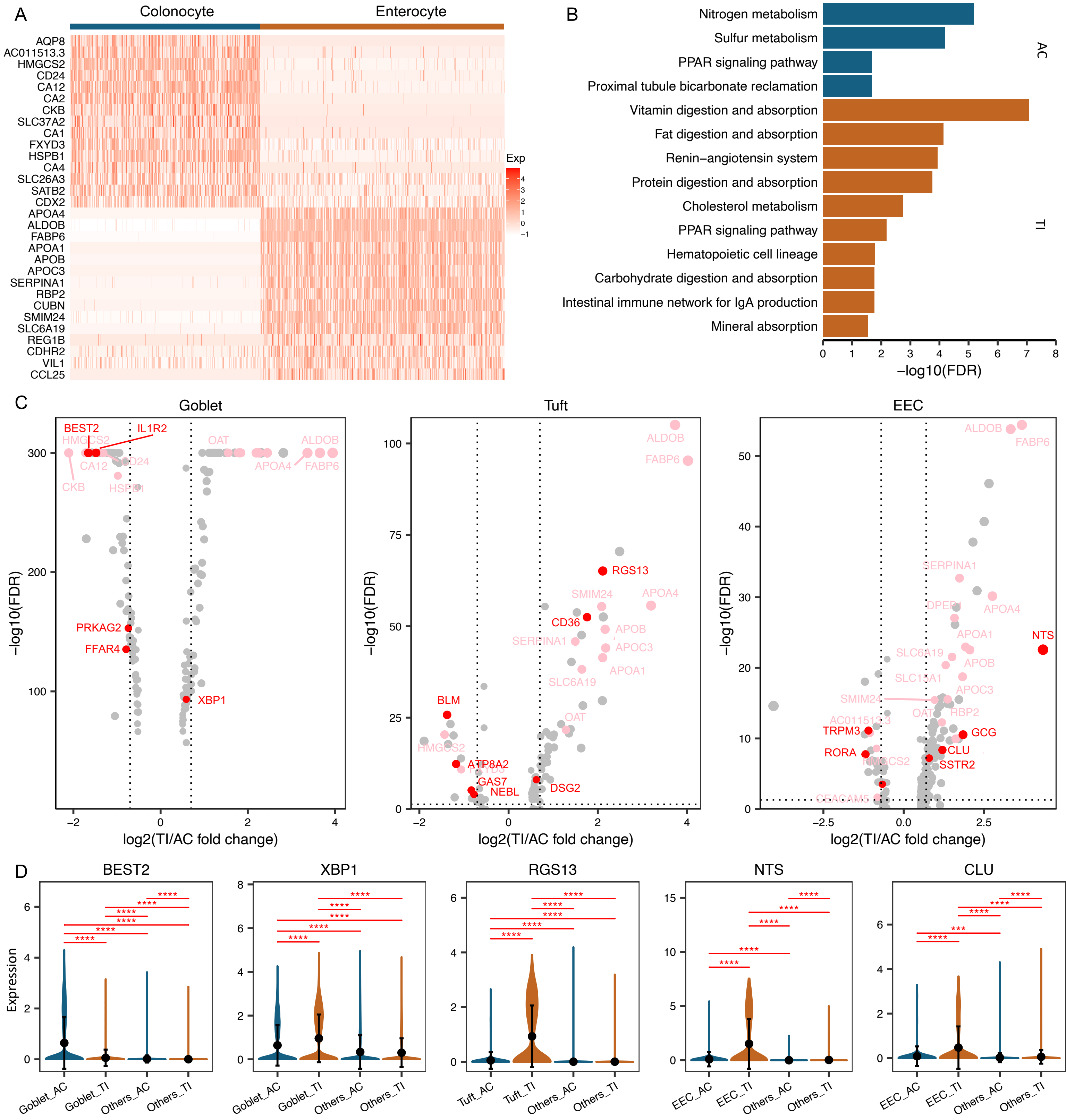


Fig. S1. A) A heatmap of the top 15 differentially expressed genes between AC colonocytes and TI enterocytes. B) Functional enrichment results on differentially expressed genes between AC colonocytes (blue) and TI enterocytes (brown). C) Volcano plots of differential results between TI goblet and AC goblet (left), between TI Tuft and AC Tuft (middle), and between TI EEC and AC EEC (right) (pink denotes region-specific genes, while red denotes genes with both region and cell-type specific); D) Violon plots of gene expression with both region and cell-type specific. AC and goblet-specific gene, BEST2, TI and goblet-specific gene, XBP1, TI and tuft-specific gene, RGS13, TI and EEC-specific gene, NTS, and TI and EEC-specific gene, CLU.

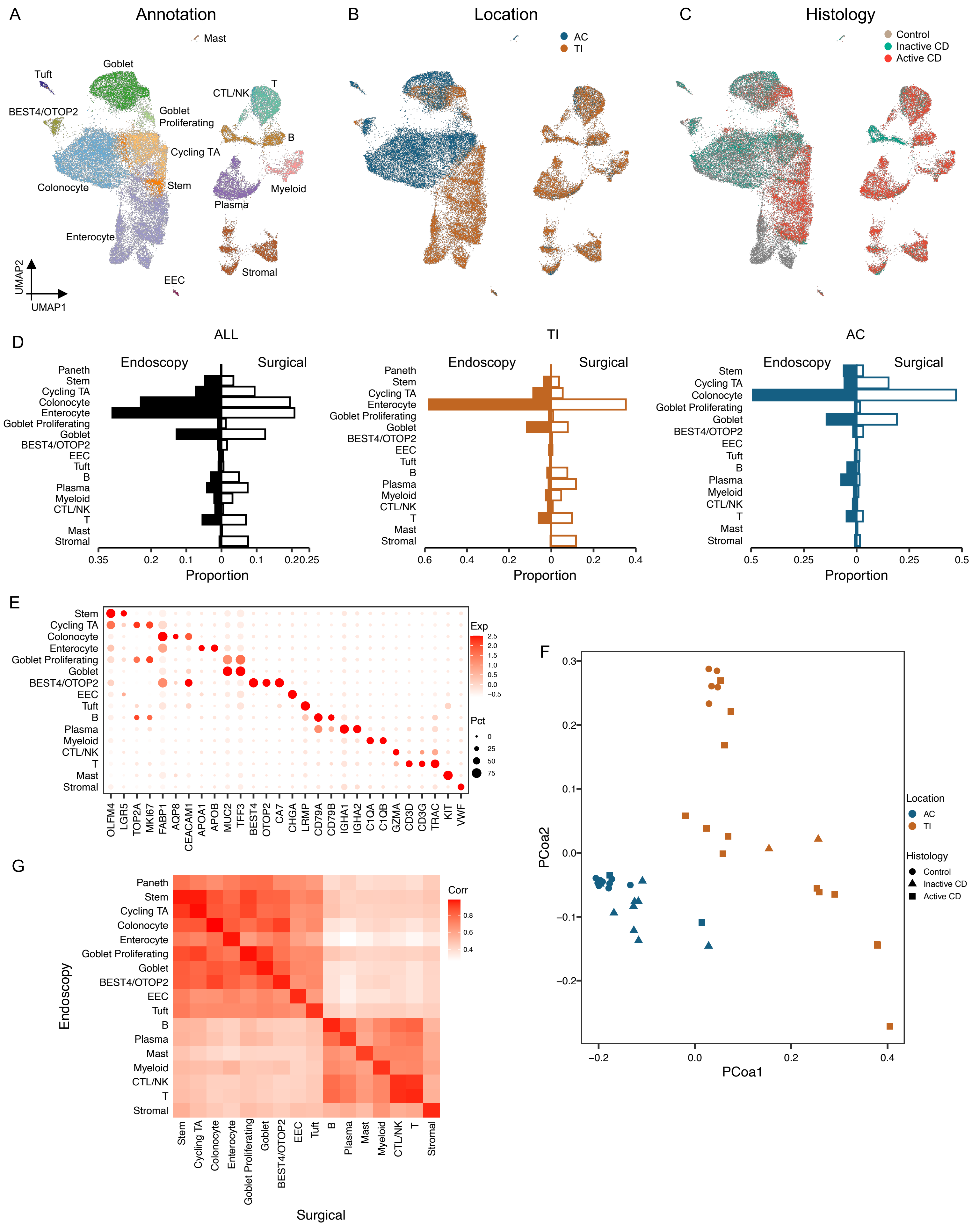


Fig. S2. A) UMAP of of 47,266 cells from surgical specimens colored by cell type; B) UMAP of 47,266 cells from surgical specimens colored by tissue origin, TI(brown) and AC (blue). C) UMAP of 47,266 cells colored by disease status, non-IBD controls (brown), inactive (green) and active CD(red). D) Comparison of cell proportions between endoscopy (filled) and surgical (unfilled) samples. E) Dotplot of marker genes in each cell type. F) MDS plot of cell compositional difference across all surgical samples. G) Similarity of cell types between endoscopy and surgical samples.

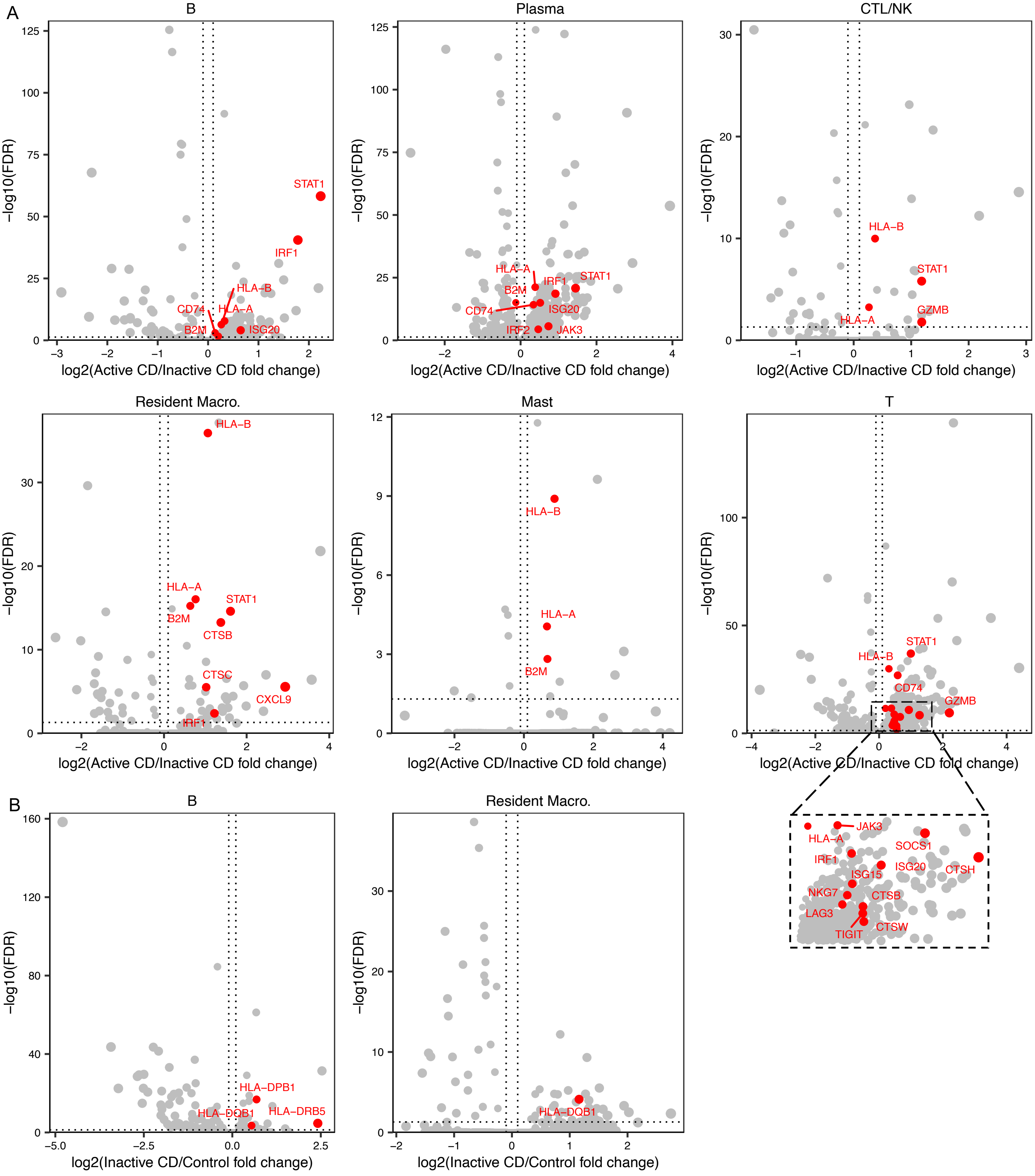


Fig. S3. A) Volcano plots of differentially expressed genes between Active and Inactive CD. B) Volcano plots of differentially expressed genes between Inactive CD and healthy controls in B and Resident Macrophages.

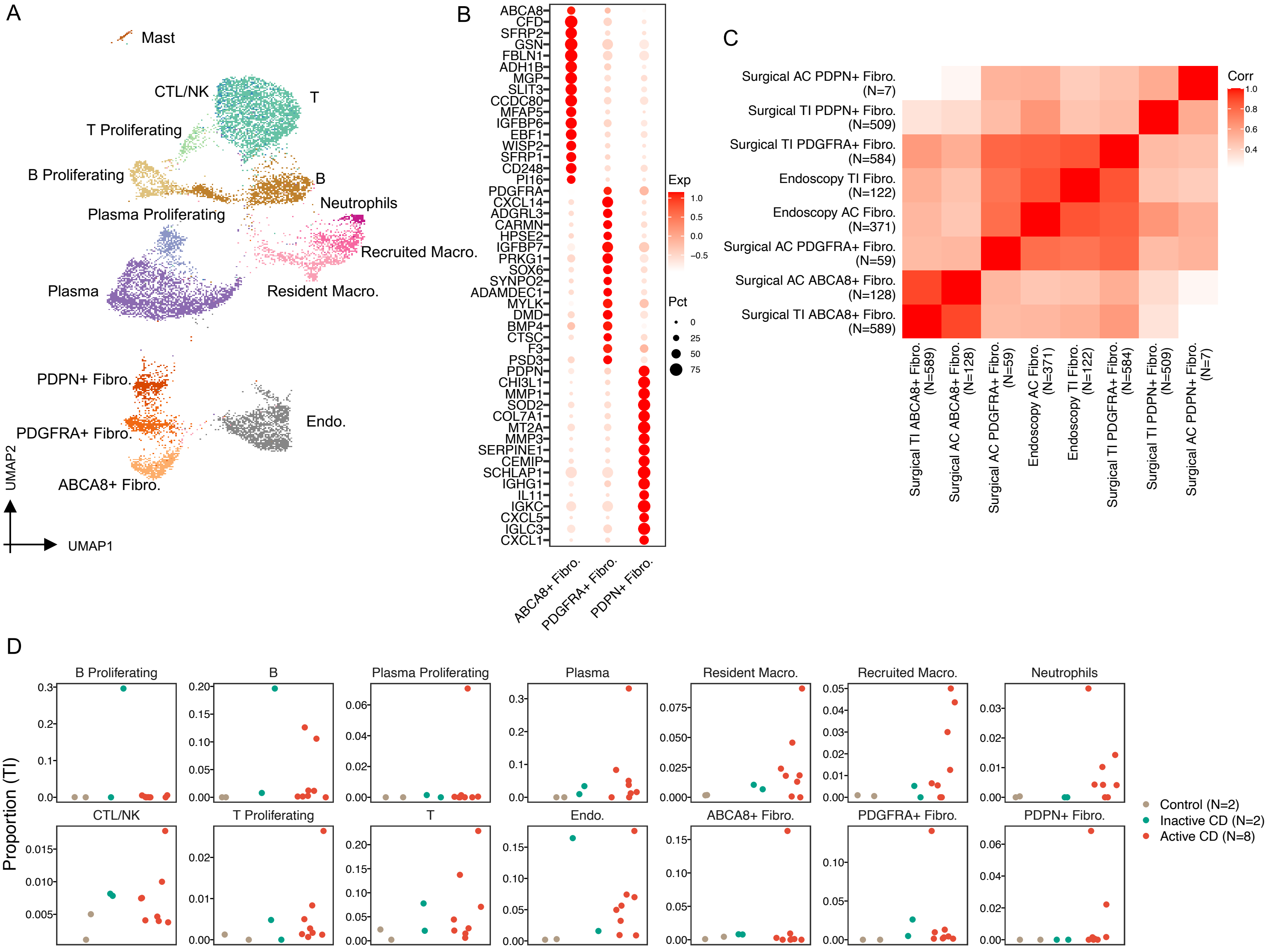


Fig. S4. A) UMAP of immune and stromal cells in surgical samples. B) Dotplot of marker genes in three fibroblast cell types. C) Similarity of fibroblast cell types detected from surgical and endoscopic samples. D) Proportional changes of immune and stromal cells from healthy controls to inactive and active CDs in surgical TI.

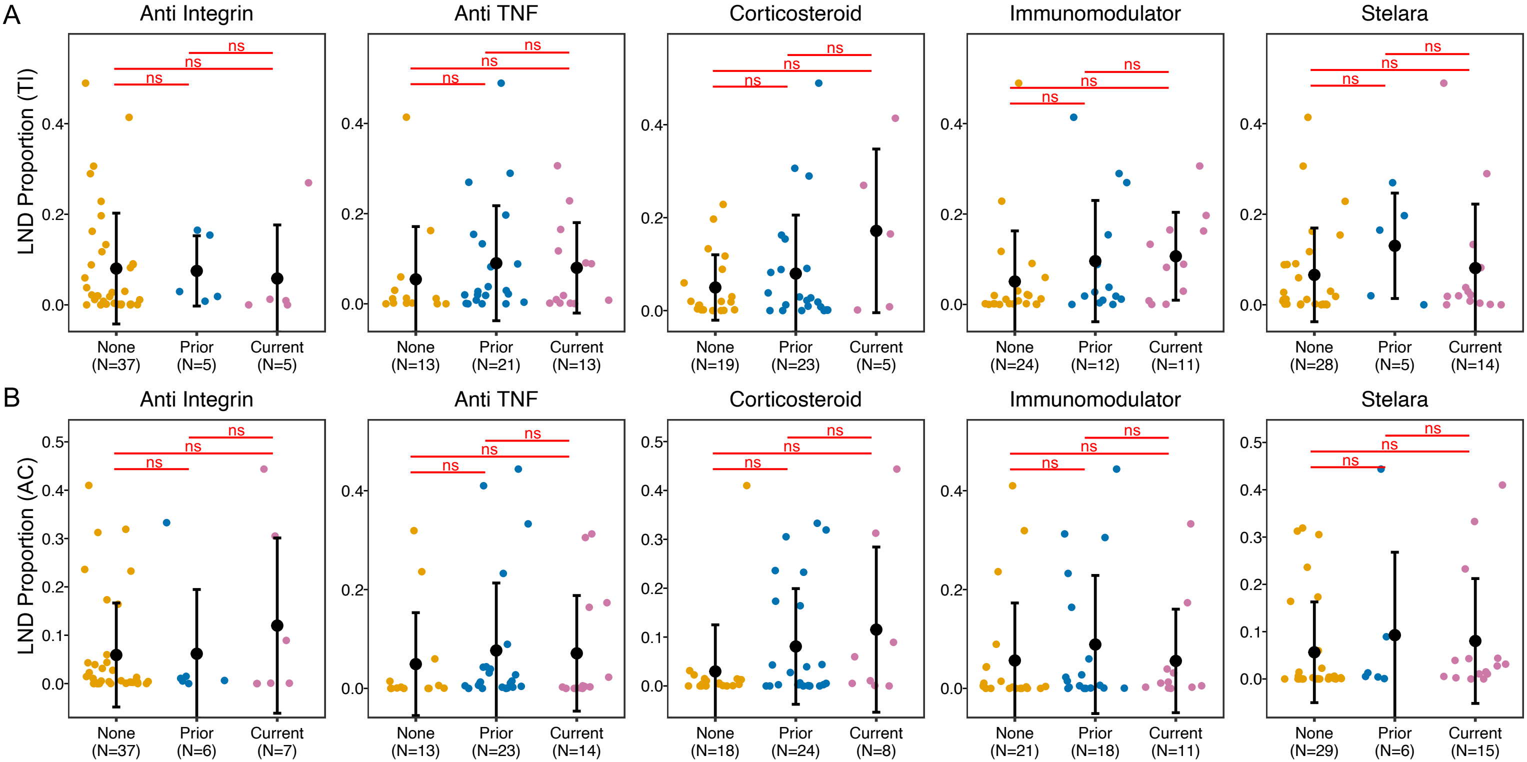


Fig. S5. LND proportional changes with different medical exposure in TI (A) and AC(B)

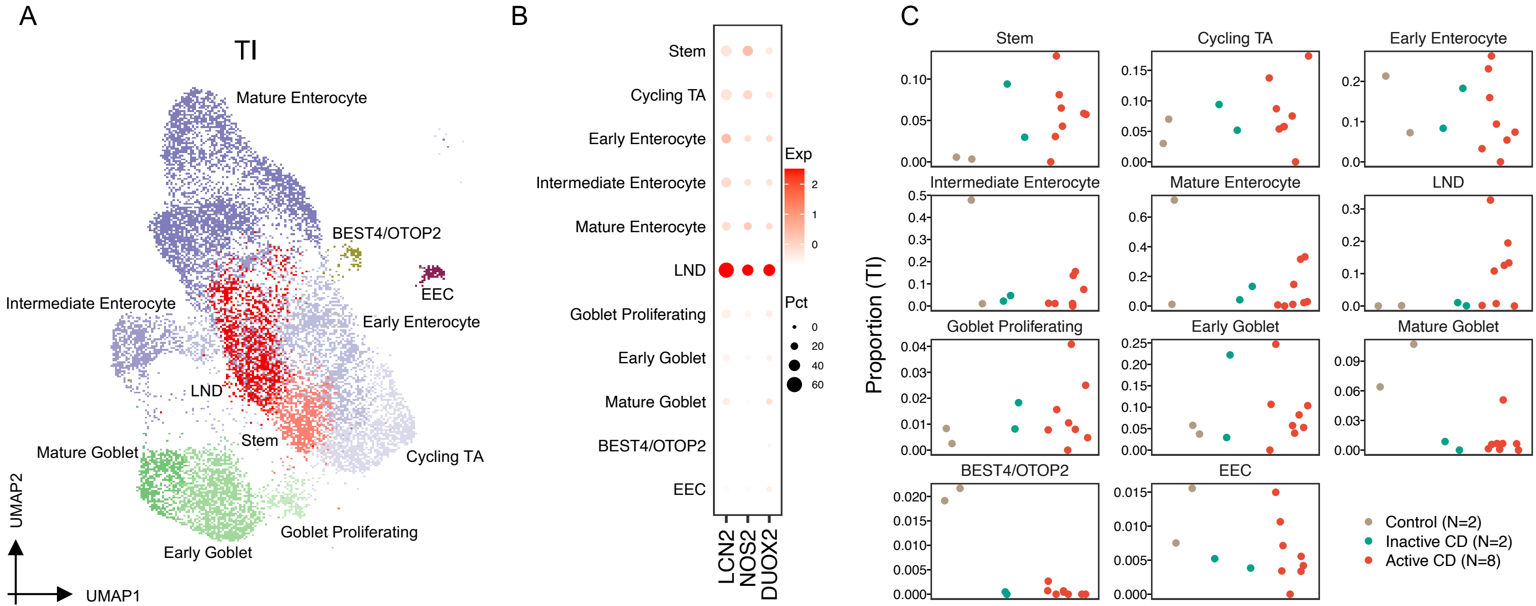


Fig. S6. A) UMAP of epithelial cells from surgical TI samples colored by cell type. B) High expression of LCN2, NOS2 and DUOX2 in the LND cluster in the surgical TI. C) Cell proportional changes of cell clusters in surgical TI from non-IBD controls, inactive and active CD.

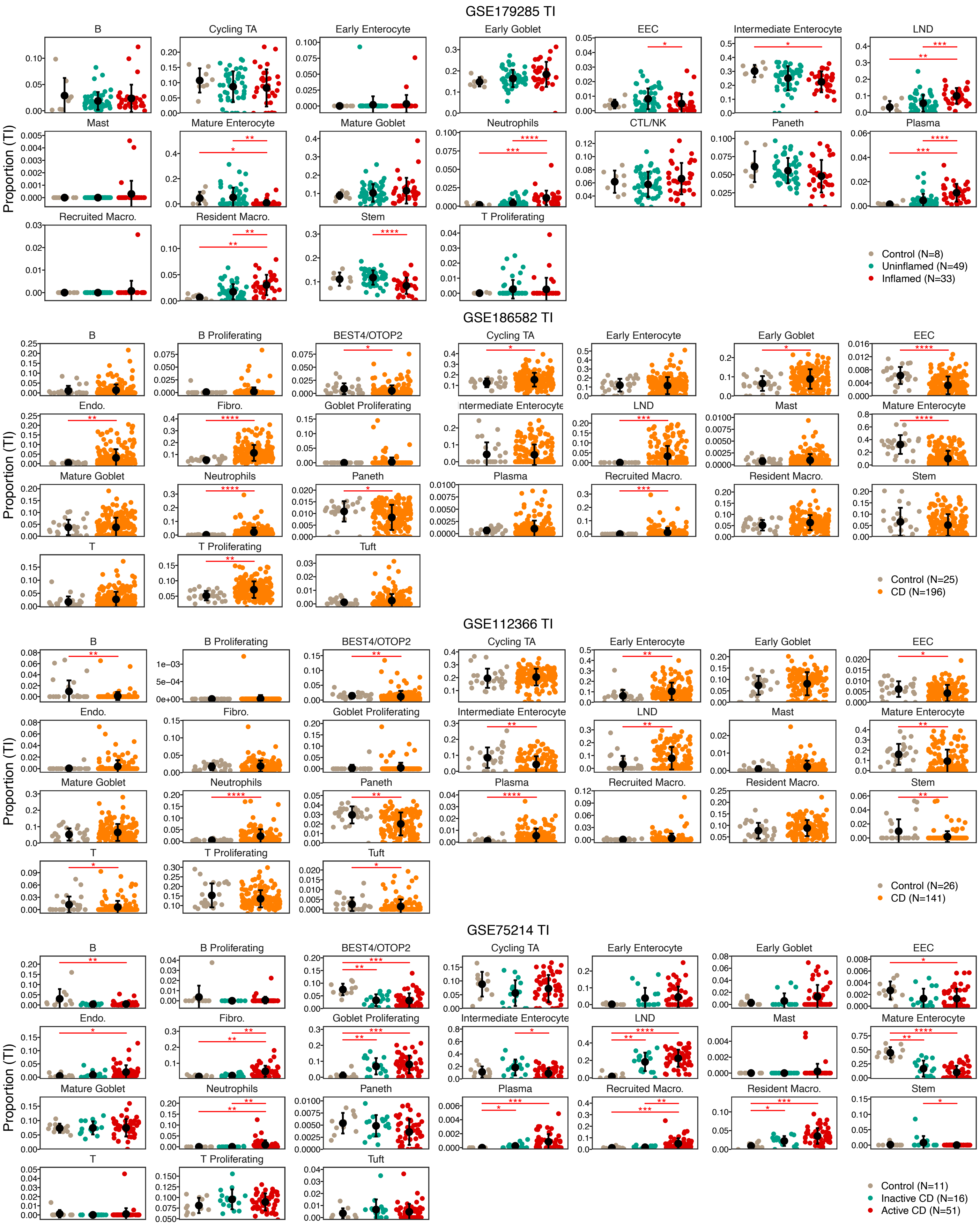


Fig. S7. Proportion changes of epithelial cells in four datasets from TI: GSE179285, GSE186582, GSE112366, and GSE75214.

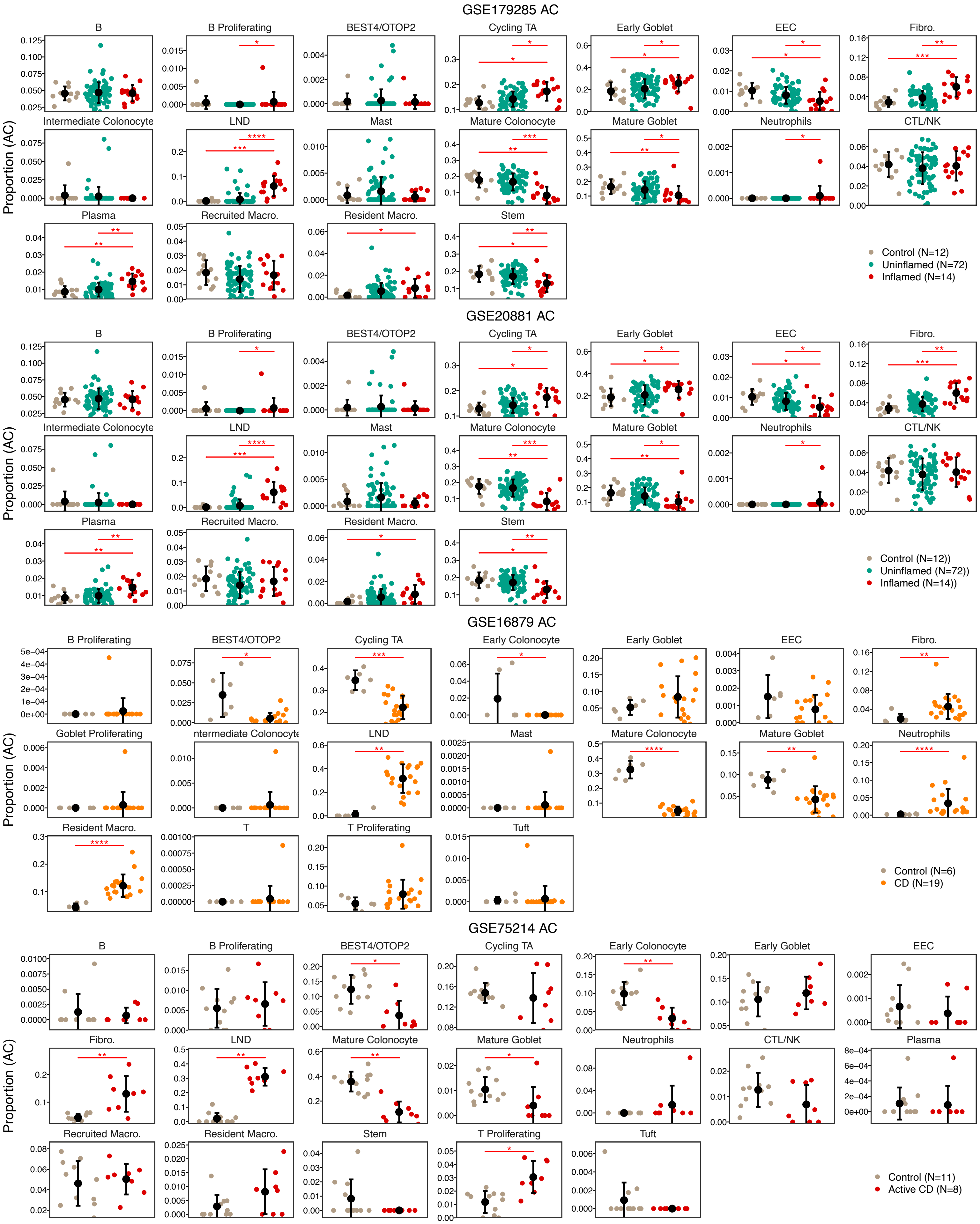


Fig. S8. Proportion changes of epithelial cells in four datasets from AC: GSE179285, GSE20881, GSE16879, and GSE75214.

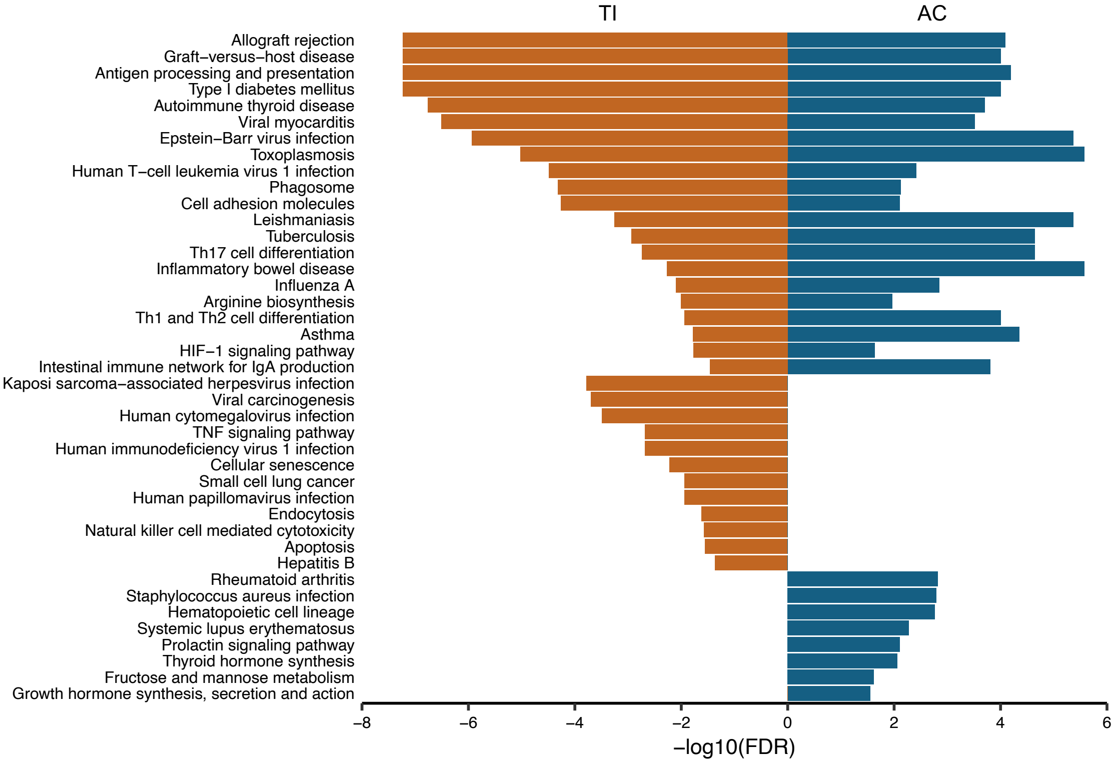


Fig. S9. Pathways enriched in the LND cluster.

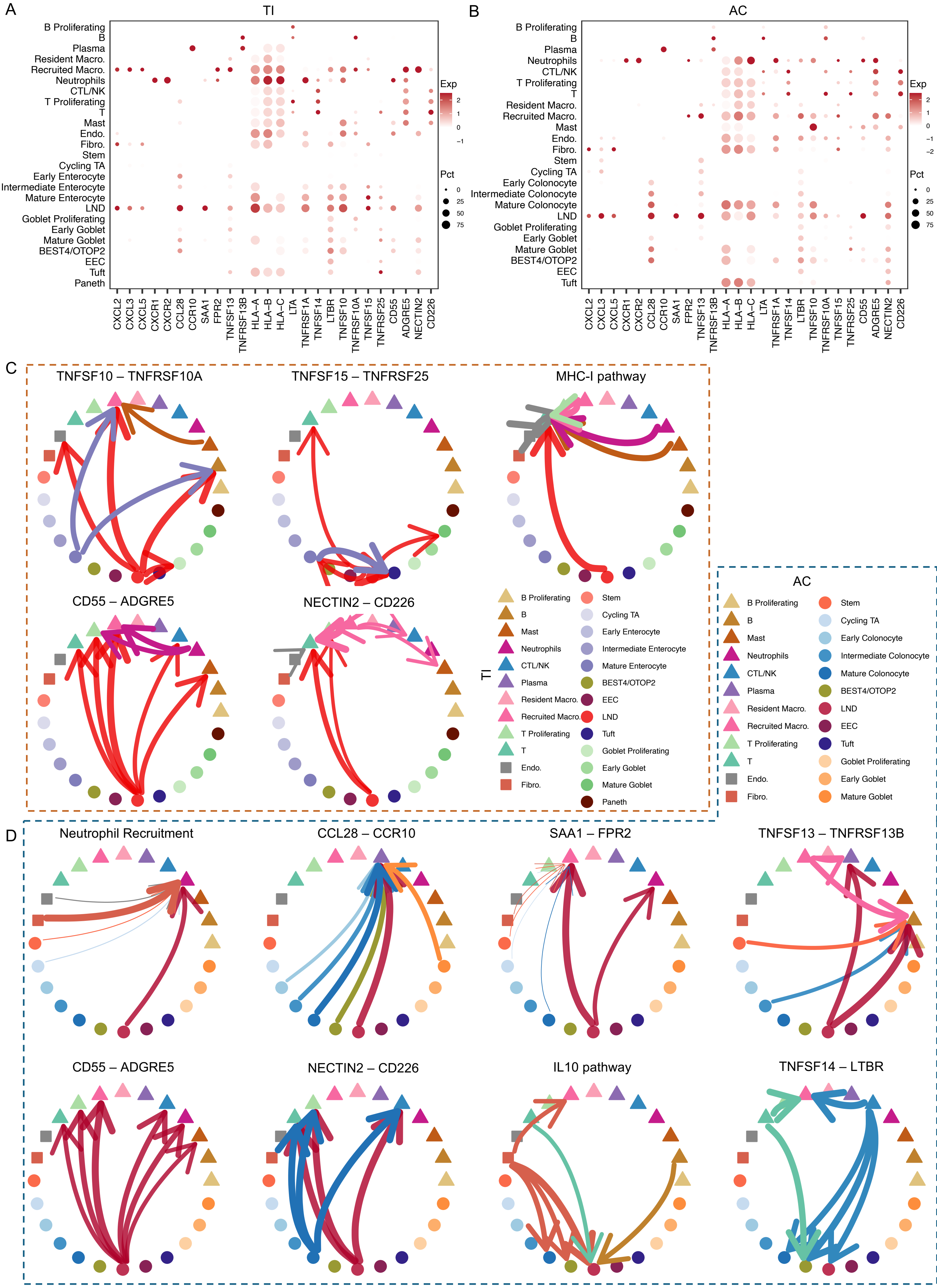


Fig. S10. Dotplots of ligands and receptors expression in different cell types in TI (A) and AC (B). Ligand-receptor interactions between LND and immune cells in TI (C) and AC (D).

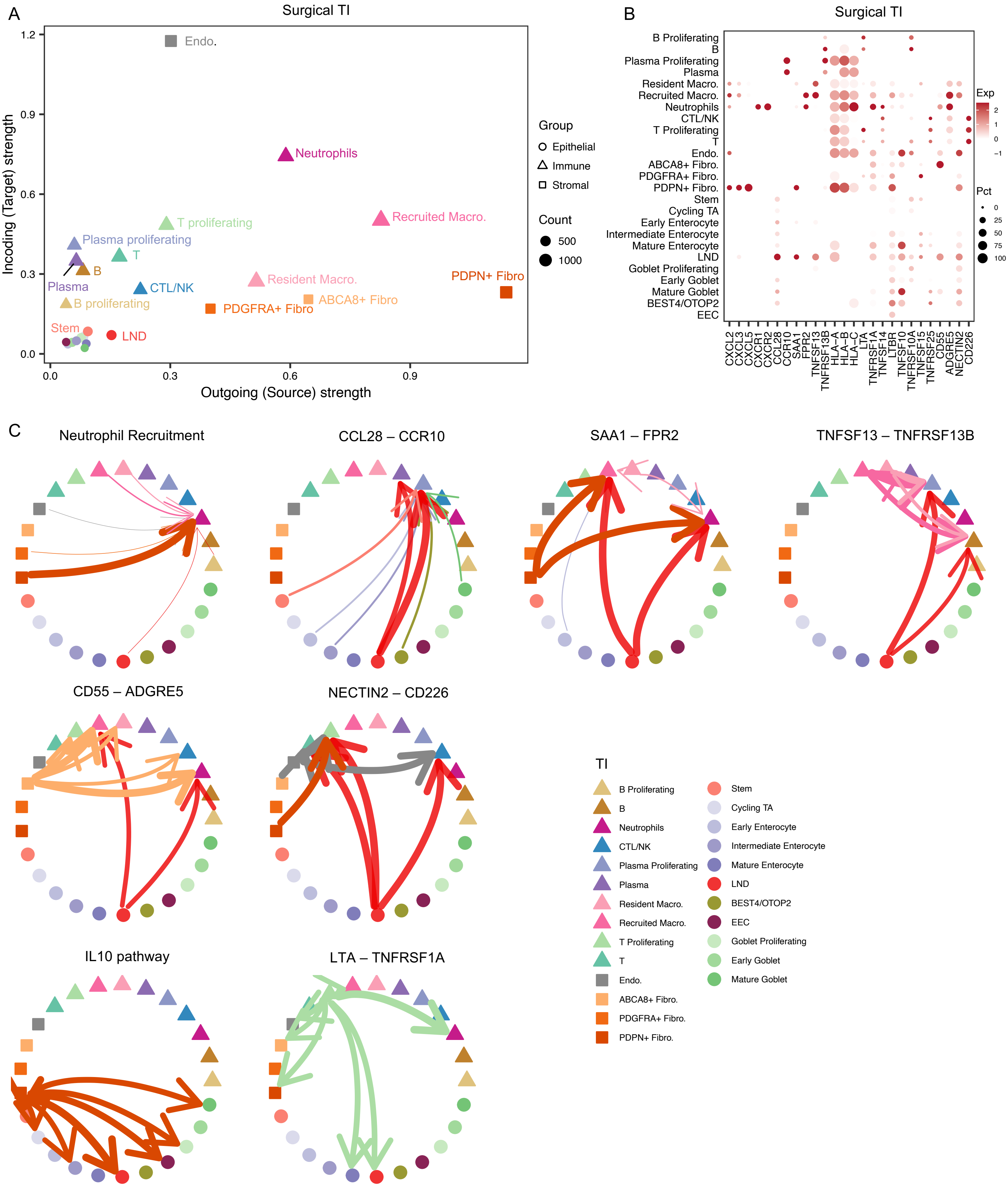


Fig. S11. A)A  scatterplot of outgoing (x-axis) and incoming (y-axis) strength of each cell type in surgical TI samples. B) DotPlot of ligands and receptors expression in different cell types in surgical TI samples. C) Ligand-receptor interactions between LND and immune cells in surgical TI samples.

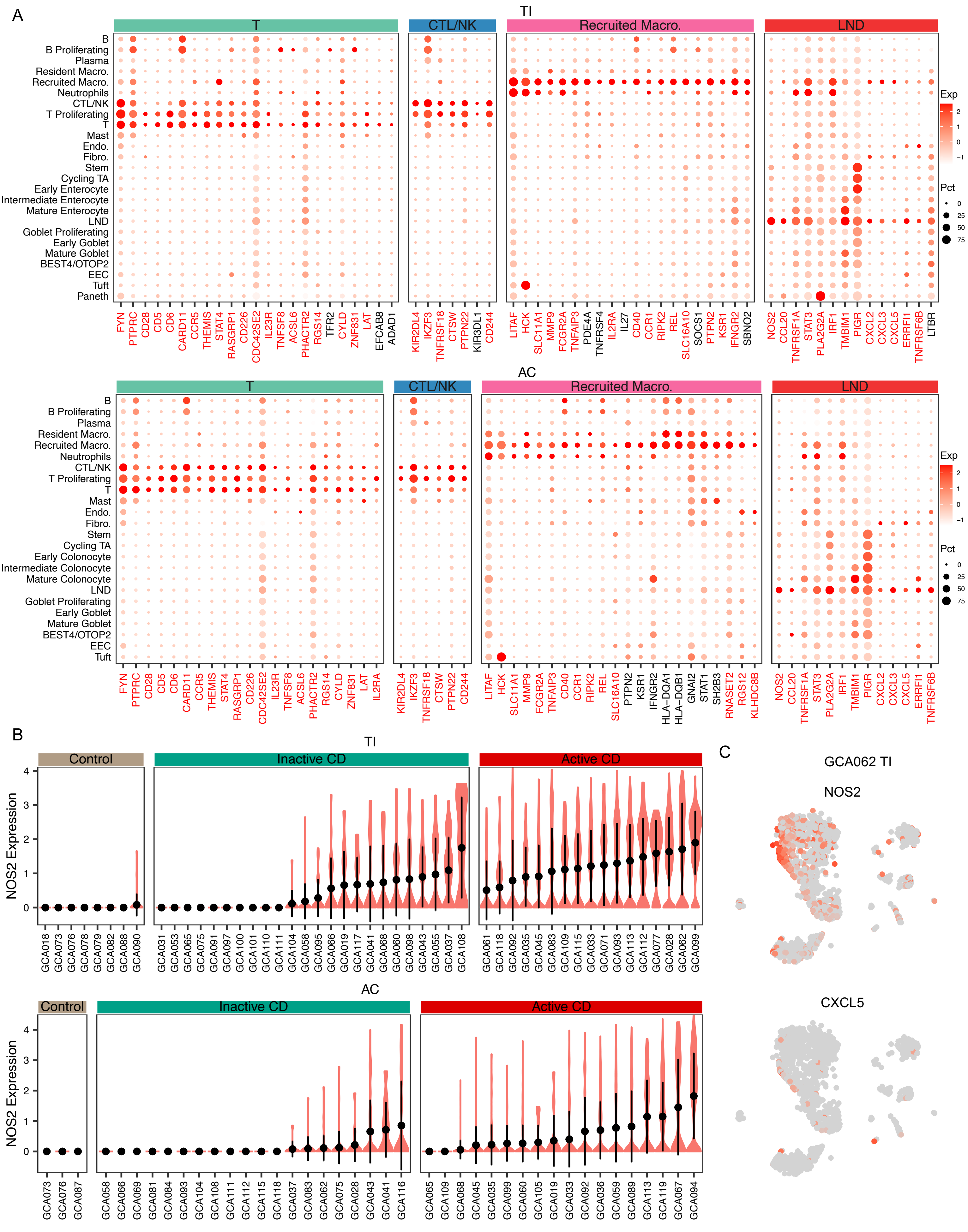


Fig. S12. A) Dotplots of expression of risk genes contributing to significant disease association of T cells, CTL/NK, recruited macrophages, and LND in TI(top) and AC (bottom). B)Expression heterogeneity of NOS2 in LND across all TI (top) and AC (bottom) samples. C) UMAP plot of NOS2 and CXCL5 expression in the TI from CD patient GCA062.

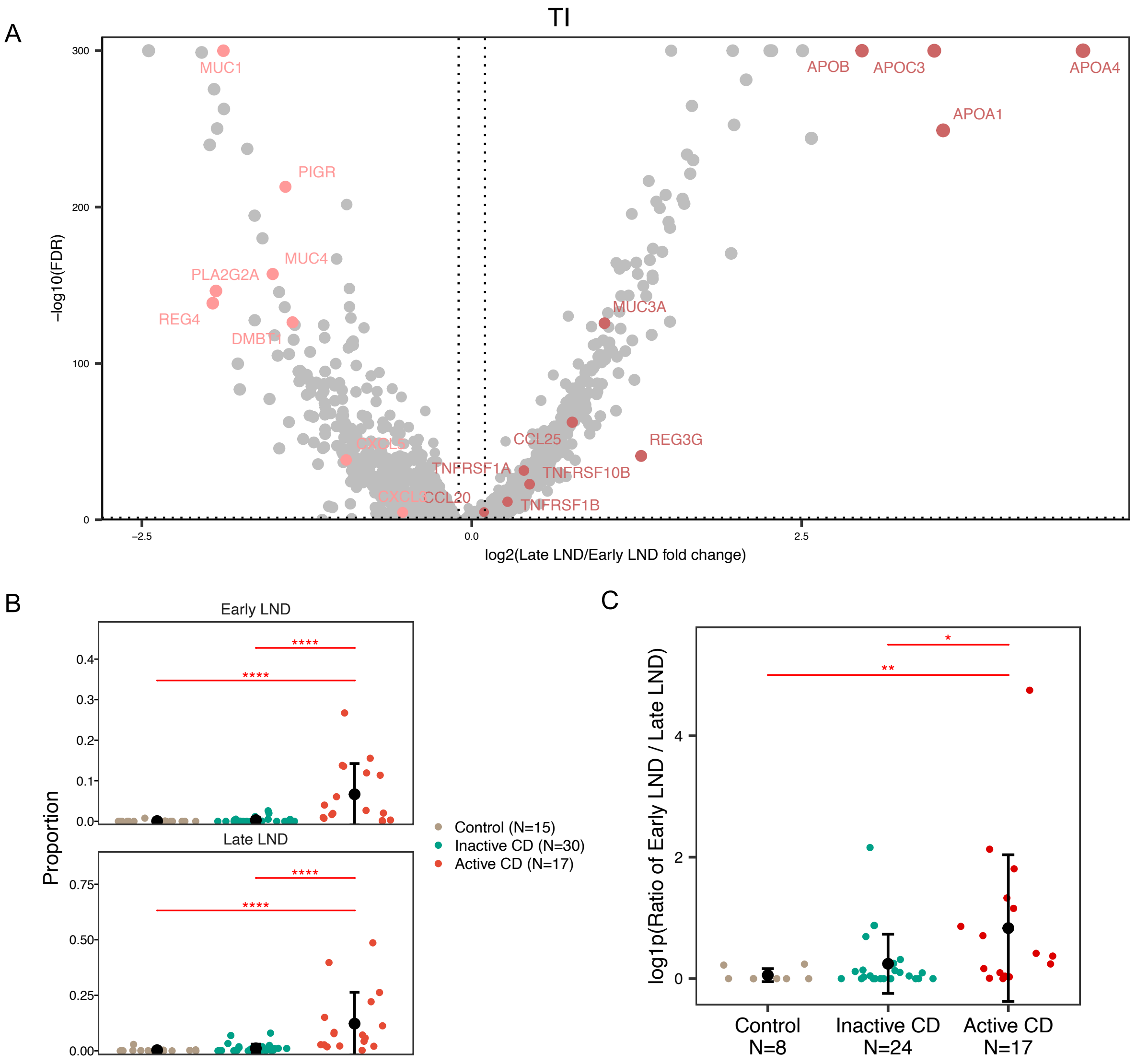


Fig. S13. A) Volcano plot of differential expression between the early and late LND in TI. B) Cell proportions of early and late LND in TI in healthy controls, inactive and active CD. C)The ratio of early to late LND in TI in healthy controls, inactive and active CD.

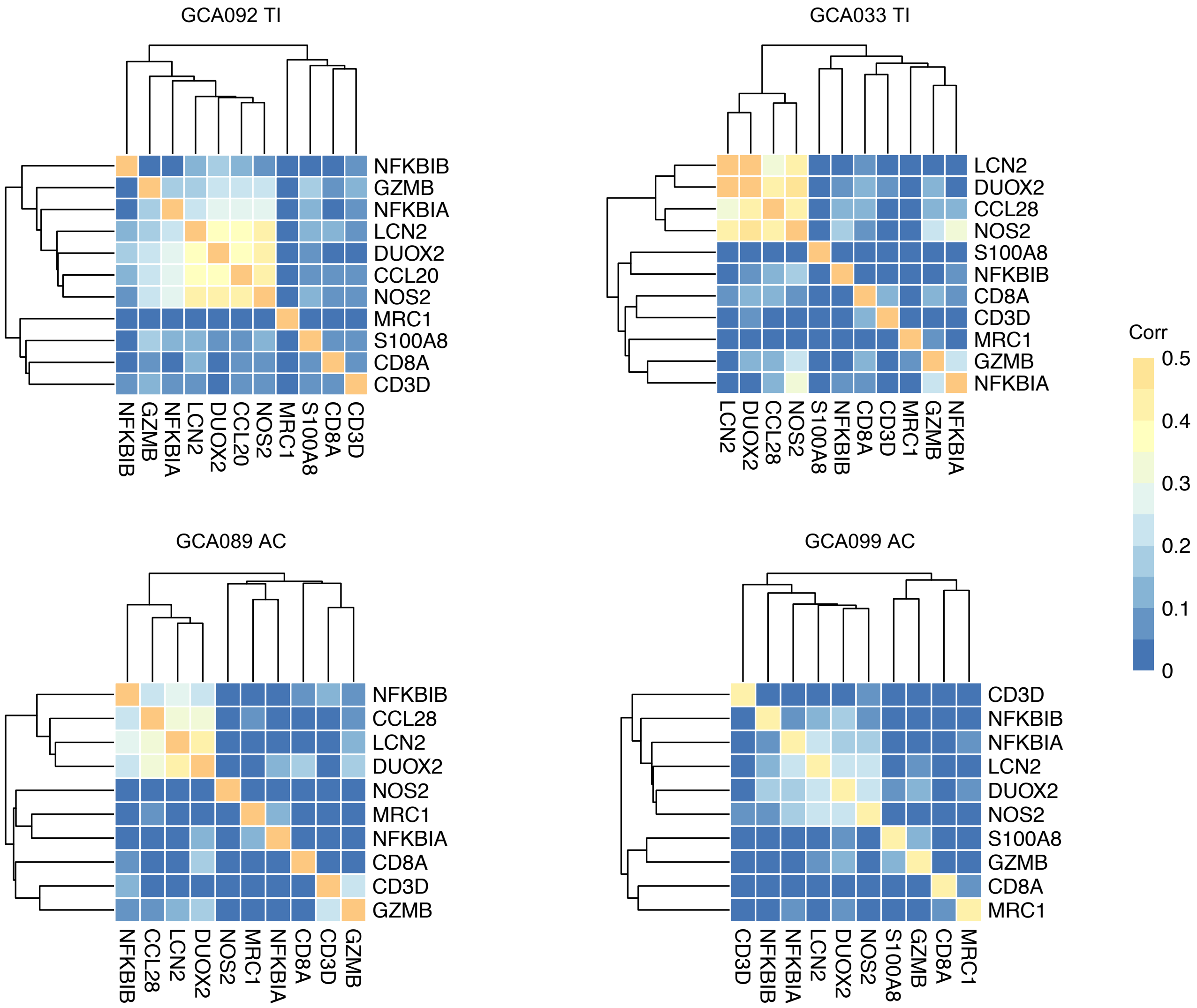


Fig. S14. Expression correlation of LND markers (LCN2,DUOX2, NOS2, CCL20 or CCL28) and immune signatures (CD3D, CD8A, GZMB, MRC1, S100A8, NFKBIA, and NFKBIB) in the four 10x Visium datasets.
